# A recent domestication event in *Penicillium biforme,* independent of the emblematic cheese mold *P. camemberti*

**DOI:** 10.64898/2026.07.27.740988

**Authors:** Samuel O’ Donnell, Gabriela Rezende, Jean-Philippe Vernadet, Alodie Snirc, Amandine Labat, Monika Coton, Elisabeth Poirier, Baris Weber, Jörg-Peter Schnitzler, Tatiana Giraud, Jeanne Ropars

**Affiliations:** Universite Paris-Saclay, CNRS, AgroParisTech, Ecologie Société Evolution, 91190 Gif-sur-Yvette, France; Universite Brest, Laboratoire Universitaire de Biodiversité et Ecologie Microbienne, F-29280 Plouzané, France; Research Unit Environmental Simulation, Helmholtz Munich, D-85764 Neuherberg, Germany; Ecosystem Physiology, Faculty of Environment and Natural Resources, University Freiburg, D-79110 Freiburg, Germany; Department of Plant Pathology, University of Wisconsin-Madison, Madison, WI 53706, USA; Quadram Institute Bioscience, Norwich Research Park, Norwich, NR4 7UQ, UK

**Keywords:** domestication, horizontal gene transfer, *Penicillium camemberti*, human-made environments, structural variants

## Abstract

Domestication of molds for cheese production has repeatedly shaped *Penicillium* fungi, most notably giving rise to the emblematic *P. camemberti* lineage, derived from *P. biforme.* Here, we identified a new *P. biforme* lineage, named *cheesy,* likely selected from *P. biforme* for food fermentation, including cheese and sausage production. This lineage exhibits evidence of a severe bottleneck, with little nucleotide polymorphism and a single mating type. The *cheesy* lineage has evolved advantageous traits for cheesemaking: compared to other *P. biforme* strains and its wild relative *P. fuscoglaucum,* it displays faster growth on cheese, galactose and lactose media, higher sporulation and germination rates on cheese, elevated lipolytic activity, enhanced inhibition capacities, and produced specific volatile organic compounds. Additionally, *P. camemberti* and *P. biforme cheesy* differ in their content of *Starship* mobile elements, acquired through horizontal transfers. These elements carry cargo genes potentially relevant for adaptation to cheese. Notably, the *cheesy* lineage has acquired a 20 kb *Starship* element (*Rattus*), nested within a much larger 160 kb *Starship (Bilge),* and carrying cargo genes with predicted functions involved in antagonistic interactions among micro-organisms.

**Significance:**

- We identified a new *Penicillium biforme* lineage, named *cheesy*, which was likely selected from *P. biforme* for food fermentation (cheese and sausage), genetically and phenotypically different from the emblematic *P. camemberti* lineage. This lineage suffered from a severe bottleneck, displaying very little nucleotide polymorphism and a single mating type.
- This newly identified lineage has evolved advantageous traits for cheesemaking; compared to other *P. biforme* strains and its wild relative *P. fuscoglaucum,* it displayed faster growth on cheese, galactose and lactose media, higher sporulation and germination rates on cheese, higher lipolytic activity, better inhibition capacities and specific volatile organic compounds.
- *Penicillium camemberti* and *P. biforme cheesy* also differed by their content in *Starship* mobile elements, acquired through horizontal transfers, and carrying cargo genes potentially relevant for adaptation to cheese.
- Notably, the *cheesy* lineage has acquired a specific 20 kb *Starship* nested within a much larger 160 kb *Starship*, and likely involved in antagonistic interactions.

## Introduction

Understanding how organisms adapt to their environment is a central question in evolutionary biology, particularly in the light of the ongoing biodiversity crisis. Domestication provides an ideal framework to study adaptation, as it involves strong and recent selection on well-defined traits, resulting in genetic and phenotypic differentiation between domesticated lineages and their wild progenitors. Fungi are valuable models to study evolution in general and domestication in particular, combining strong experimental assets with a remarkable diversity (Gladieux et al., 2014). They typically have small genomes, with easy access to the haploid phase, are experimentally tractable and many species have undergone domestication. Many fungi are used in the production of fermented foods, including wine, bread, sake, cheese, dried sausage, soja sauce and tempeh, as well as for the production of enzymes (e.g. lactase), bioactive compounds such as mycophenolic acid for immunosuppressant medication and antibiotics, including penicillin. Yet, the mechanisms involved in the rapid adaptation occurring during microbe domestication remain poorly understood.

Horizontal gene transfer (HGT), i.e. the non-sexual transfer of genetic material between species, has long been recognized as a major driver of prokaryotic evolution and adaptation (Arnold et al., 2022), and is increasingly reported in eukaryotic microorganisms (Etten & Bhattacharya, 2020). Newly acquired elements can confer significant fitness benefits, such as increased efficiency of nutrient uptake or competitive advantages, through increased growth rate or toxin production. Horizontally transferred regions (HTRs) between cheese *Penicillium* species have been documented (Cheeseman et al., 2014; Ropars et al., 2015), likely involved in lactose metabolism and competition. These HTRs were retroactively identified as *Starships,* a recently described superfamily of giant transposable elements widespread across fungal lineages (Urquhart et al., 2024)*. Starships* have been suggested to facilitate rapid adaptation by shuttling cargo genes between strains and across species. *Starships* have been found in fungal human and plant pathogens (Bucknell et al., 2025; Gluck-Thaler et al., 2025; Hartmann et al., 2025; Peck et al., 2024; Sato et al., 2025), as well as in *Penicillium* and *Aspergillus* molds domesticated for the fermentation of food, including cheese but also dry-cured meat (Lo et al., 2023; O’Donnell et al., 2025). These *Starships* carried cargo genes with functions relevant for adaptation to their respective environments. *Starships* are characterized by the presence of a tyrosine recombinase at one extremity, a.k.a. the Captain, which has been shown to be essential for *Starship* movement (Urquhart et al., 2023) and is used to detect *Starships* in genomes (Gluck-Thaler & Vogan, 2024; O’Donnell et al., 2025)*. Starships* are more abundant in domesticated fungal lineages than in their wild counterparts, and are particularly prevalent in cheese-associated molds (O’Donnell et al., 2025).

Among domesticated fungi, the most emblematic ones may be the molds used for maturing cheeses. These domesticated cheese fungi are expected to display specific traits beneficial for cheese making, such as faster growth in cheese at cave temperatures compared to wild relatives, allowing speeding up maturation processes, and reduced proteolytic activities, these negatively impacting cheese flavors when too rapid. In addition, traits affecting cheese aroma may also have been targets of domestication, as fungal volatile organic compounds (VOCs) contribute to the metabolic signature of ripening cheeses (Bertuzzi et al., 2018; Molimard & Spinnler, 1996). In contrast, the ability to grow in harsh conditions, e.g. with limited nutrients and high competition levels, may have been lost in domesticated cheese fungi due to relaxed selection (Ropars et al., 2020), as often reported for unused traits in human-made environments in domesticated organisms. This can affect traits related to nutrient acquisition or toxin production involved in competition. They may also have suffered from bottlenecks during domestication.

One of the most iconic cheese molds is *Penicillium camemberti.* This species is commonly used to ripen soft cheeses like Camembert, giving them their characteristic white, fluffy rind. *Penicillium camemberti* derives from a mutant line selected in 1898 from its blue-grey wild progenitor, called at the time *P. commune* (Delfosse, 2008). Genomic data showed that *P. commune* was not monophyletic but instead comprised strains belonging to either *P. fuscoglaucum* or *P. biforme* (Ropars et al., 2020); *P. commune* is therefore no longer a valid taxonomic name. Genome analyses further confirmed that *P. camemberti* was a clonal lineage, with very low diversity, selected for its white and fluffy aspect from a *P. biforme* population (Ropars et al., 2020). Two varieties are differentiated within *P. camemberti*, var. *camemberti* and var. *caseifulvum,* which display markedly different phenotypes, while both exhibit beneficial traits for cheese making, that are lacking in *P. biforme* and their wild relative *P. fuscoglaucum*. The variety *camemberti* of *P. camemberti* is whiter and fluffier, and inhibits more efficiently fungal competitors. The variety *caseifulvum* is more grey-green, has reduced aerial hyphae and faster growth on cheese; it has lost its ability to produce cyclopiazonic acid, a mycotoxin, due to a frameshift in one of the genes of the biosynthetic cluster (Ropars et al., 2020). Similarly, the domesticated lineages of the blue cheese mold, *P. roqueforti,* display traits beneficial for cheese making, differentiated from non-cheese populations, with contrasting traits between cheese lineages (Crequer et al., 2023; Dumas et al., 2020), in particular faster growth on cheese and higher levels and diversity of volatile organic compounds. Some domesticated *P. roqueforti* lineages have lost their ability to produce mycophenolic acid, another mycotoxin (Crequer et al., 2024; Gillot et al., 2017). Domesticated *P. camemberti* and *P. roqueforti* populations show very low genetic diversity (Crequer et al., 2023; Dumas et al., 2020; Ropars et al., 2020) and share most of their *Starships* (Cheeseman et al., 2014; Ropars et al., 2015), some of which are also shared with the *Penicillium* molds used for maturing dry-cured meat (Lo et al., 2023).

*Penicillium biforme* also occurs naturally on a variety of cheeses and is easily recognizable by its distinctive light blue-grey color, but its diversity, evolutionary history and *Starship* diversity have hardly been studied due to limited sampling. It is particularly common in artisanal cheeses, such as hard, semi-hard, blue-veined and fresh varieties, where no fungal cultures are intentionally inoculated (Irlinger et al., 2024; Penland et al., 2021). As a result, *P. biforme* typically appears as scattered spots on cheeses, unlike the inoculated *P. camemberti,* which forms a uniform, immaculate layer on soft cheeses like Camembert or Brie. While industrial cheese production may occasionally include *P. biforme* as an inoculated culture, it is added in more limited amounts, preventing it from ever becoming the dominant mold.

Here, we aimed to investigate further the population structure and domestication history of *P. biforme* and *P. camemberti* for cheese making, including the potential role of *Starships*. For these goals, we have expanded our strain collection to include a total of 75 *P. biforme* strains from cheeses and other environments, as well as 22 *P. camemberti* strains. Genomes were sequenced for all strains, including two new long-read assemblies, and multiple phenotypic traits measured. Our general aim was to assess whether additional lineages have been domesticated from *P. biforme*, in addition to *P. camemberti,* i.e., whether some lineages were specific to cheeses, with genetic and phenotypic differentiation from populations originating from other environments, and what role *Starships* play in adaptation to cheese. To reach this goal, we investigated i) the genetic diversity and population structure in *P. biforme*, ii) the *Starship* presence/absence polymorphism and their potential role in adaptation, and iii) the phenotypic differences between identified populations in terms of fluffiness, radial growth on different media, on different carbon and metal sources, vertical growth, germination, sporulation, lipolytic and proteolytic activities, metabolite and volatile organic compounds (VOCs) emission, as well as toxin production.

## Results

### *A new domesticated lineage in*P. biforme*, separate from* P. camemberti

Our dataset included 97 haploid strains isolated from different cheeses (semi-hard, mold-ripened, smeared and fresh soft cheeses) or from non-cheese environments (e.g. hay, lacto-fermented cucumber, dried sausage). Among them, 75 strains belonged to *P. biforme* and 22 strains to *P. camemberti,* including 10 var. *caseifulvum* and 12 var. *camemberti*. We generated two new long-read genome assemblies, of the ESE00418 and ESE01543 *P. biforme* strains. In total, 417,650 SNPs (single nucleotide polymorphisms) were identified across the 97 strains by mapping against the ESE01543 reference genome.

The splitstree clearly separated the two *P. camemberti* varieties, and also a new clonal lineage, containing 18 food strains (mostly cheese), which we named *cheesy* (Fig. 1A). This new lineage was also separated from the other *P. biforme* strains in the admixture and PCA analyses, while the two *P. camemberti* varieties were not differentiated in these analyses (Figs. 1B, 1C, SXk2-8). The *cheesy* population contained strains isolated from dried sausages and different kinds of cheeses, but only industrial cheeses, from different countries, e.g. blue veined cheese, brie, Harbison blue, and Ossau Iraty. The *cheesy* lineage displayed low genetic diversity (π = 3.72e-05, theta = 4.29e-05), even less than the clonal domesticated *P. camemberti* var. *camemberti* (π = 1.49e-05, theta = 1.92e-05). All 18 strains shared the same mating type, MAT1-2. In contrast, among the remaining *P. biforme* strains, 36.8% carried the MAT1-1 haplotype. These results altogether suggest that the *cheesy* lineage is a commercial clonal culture of a single strain, as for *P. camemberti*.

**Fig. 1.**
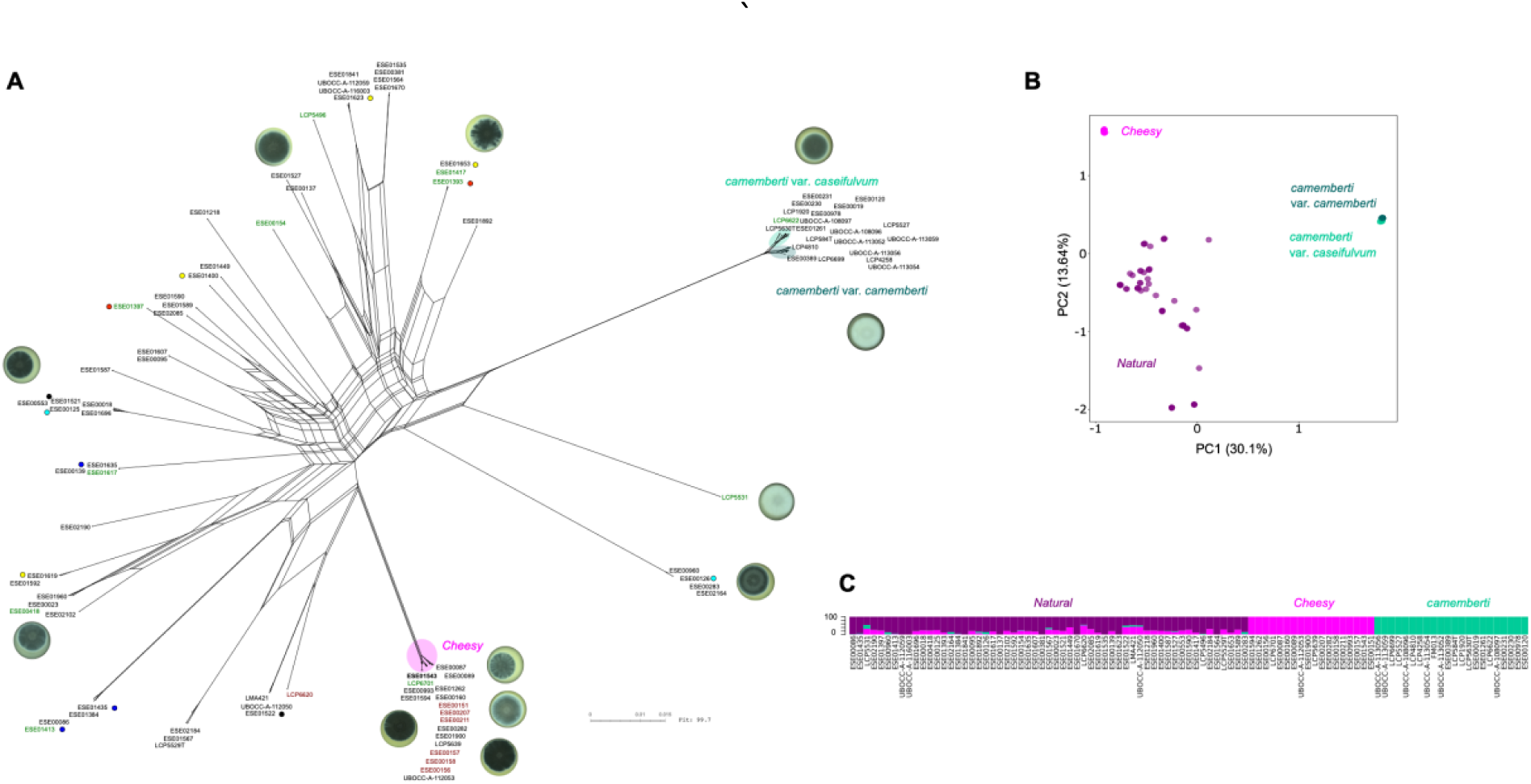
Population structure of *Penicillium biforme*. A) Neighbor-net performed with a dataset of 417,650 SNPs, including 97 taxa belonging to *Penicillium biforme* and the two varieties of *P. camemberti.* Same color circles next to strain IDs indicate strains coming from the same artisanal cheese producer. Strain IDs are colored according to their origin: black, cheese; red, dried sausage; green, other environments than food. The reference genome for all analyses, ESE01543, is highlighted in bold. B) PCA (principal component analysis) showing three genetic clusters, corresponding to *P. camemberti, cheesy* and *natural P. biforme.* C) Barplots of the NGSadmix analysis at K = 3.

The remainder of *P. biforme* corresponds to a natural population, encompassing strains isolated from non-inoculated artisanal cheeses and other environments than cheese, such as hay. *Penicillium biforme* is thus likely naturally present in the environment, either contaminating the milk during cheese making, or directly the rind later, in the ripening caves. The presence of genetically different strains in cheeses of the same artisanal cheese producer (colored circles, Fig. 1A) is consistent with a natural contamination of cheeses from the environment, instead of a commercially inoculated culture. We therefore named this population “*natural”*; its genetic diversity was relatively high (π = 1.71e-03; theta = 9.82e-04), similar to what was found in some of the non-clonal domesticated populations of the cheese-making fungus *Geotrichum candidum* (populations cheese_1 and cheese_3, (Bennetot et al., 2023)). The two mating-type haplotypes, typical of ascomycete fungi, were detected in the *natural P. biforme* population, and reticulations were observed in the Splitstree (Fig. 1A), suggesting that sexual reproduction occurs. No subdivision was found according to the environment of collection (Fig. 1A). Genetic differentiation was high between *P. camemberti, P. biforme cheesy* and *P. biforme natural* (Table S1).

### *Abundant* Starships *fuel genomic variability in* Penicillium biforme *despite low nucleotide diversity*

By using as proxy the ‘Captain’, i.e. the DUF3435 domain, we detected striking differences in terms of *Starship* content between *P. biforme, P. camemberti* and *P. fuscoglaucum* populations. We also observed a presence/absence polymorphism within populations, even within the clonal lineages with very low SNP diversity (Figs. 2 and 3, Table 1). Variation was nevertheless lower within than between populations, with higher *Starship* copy numbers in *P. camemberti* var. *caseifulvum, P. biforme cheesy* and *natural* populations than in *P. camemberti* var. *camemberti* or the wild relative *P. fuscoglaucum* (Figs. 2 and 3, Table 1).

**Fig. 2.**
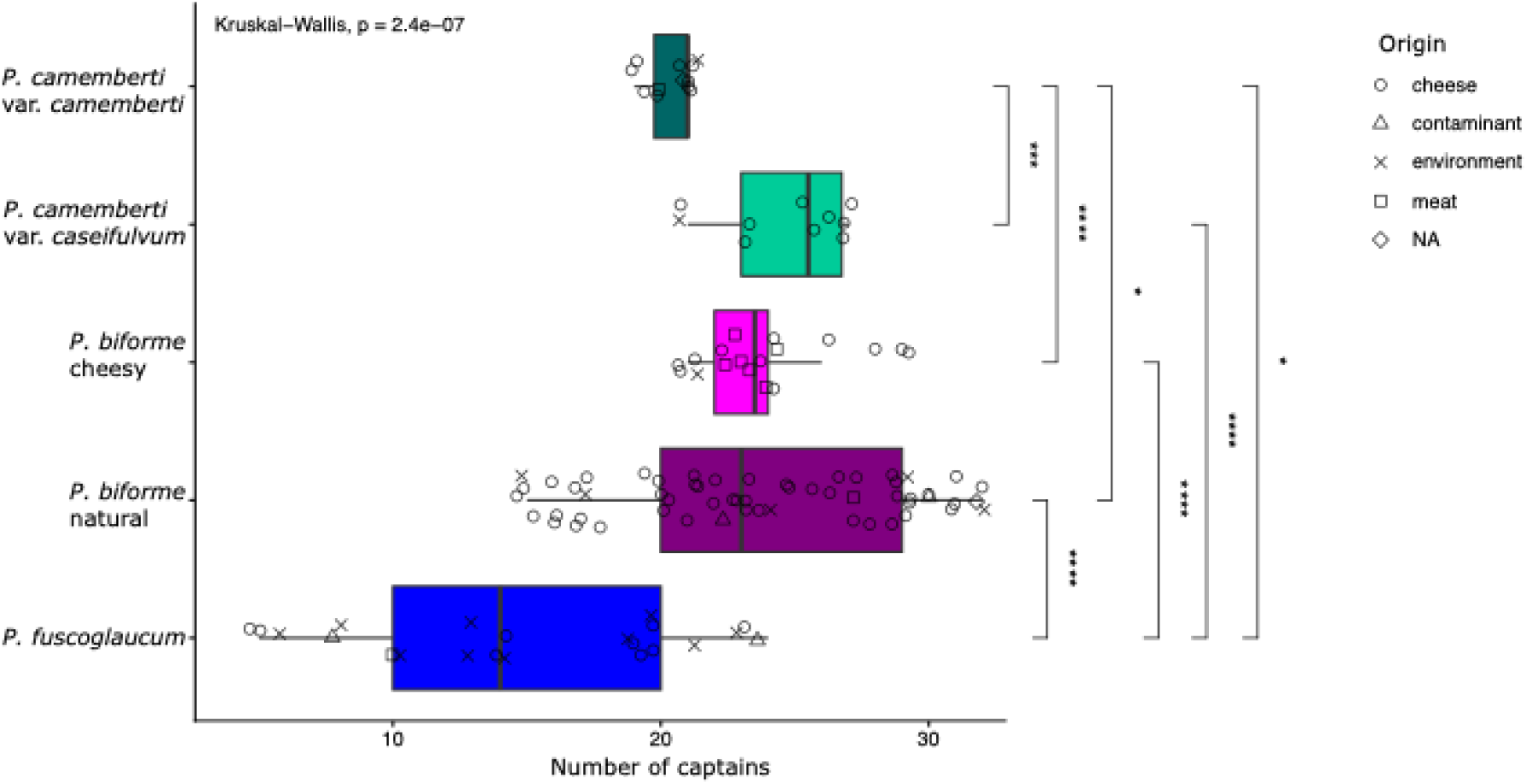
Number of captains in the various strains, indicating the *Starships* content in *Penicillium camemberti* var. *caseifulvum,* var. *camemberti, P. biforme* and their wild relative *P. fuscoglaucum*. Points represent strains, classified by species/population along the Y axis, and the shapes of the points indicate the origin of the strains (cheese, meat, contaminant, other environments, or NA for non-assigned). Only significant comparisons are shown (Wilcoxon test, * = p <= 0.05; ** = p <= 0.01; *** = p <= 0.001, Table S2).

**Fig. 3.**
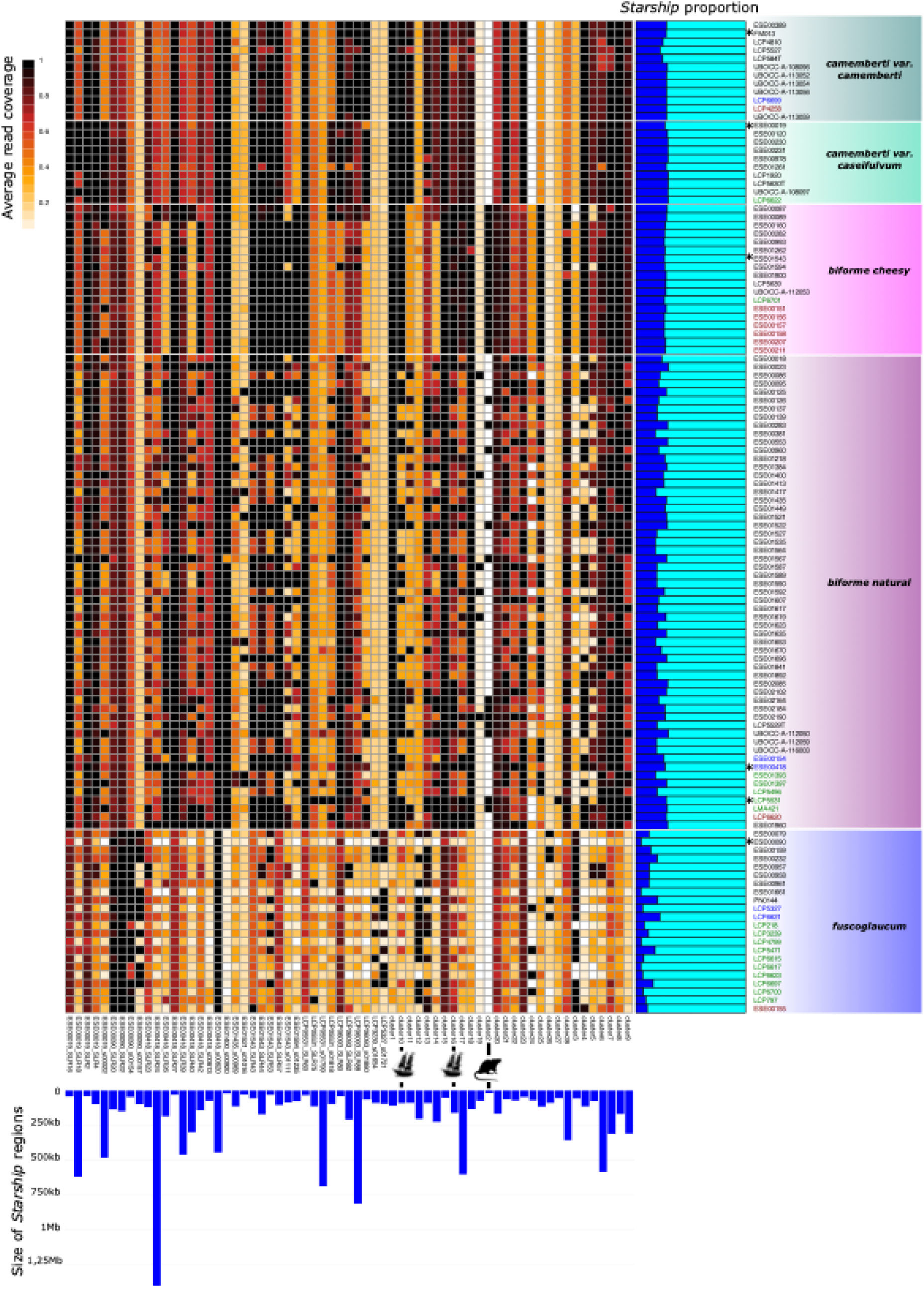
Presence/absence of *Starship-*like regions in genomes of *Penicillium camemberti, P. biforme* and *P. fuscoglaucum* strains (left panel) and proportion of *Starship* material in each genome (right panel). The presence/absence of *Starship-*like regions was inferred by mapping on long-read assemblies only. Left panel: Heatmap showing the average read coverage of each *Starship* element (column) in all strains (rows), white representing the complete absence of the *Starship* and black the complete presence (entire *Starship*)*. Starships* found in different strains are clustered together (named clusters 1 to 28) based on k-mer similarity (Jaccard similarity index) with a minimum of 30%; clusters are defined by default using the Markov Cluster Algorithm (MCL). Bottom: barplots representing the size of each *Starship*. Right panel: For each strain is plotted the percentage of the maximum cumulative length of all identified *Starship*-like regions (SLRs), as recovered by read mapping. A SLR was considered completely present when at least 50% of its length was covered by sequencing reads, and its full length was then included in the total. Strain names are colored according to their environment of origin: black for cheese, red for sausage, blue for food contaminants, green for other environments than food. Strains with available long-read assemblies are marked with an asterisk.

We used high-quality genome assemblies to confirm the variability in *Starship* content between and within populations. Across the long-read genome assemblies of six strains, representative of the different populations, we identified 25 *Starship-*like regions (O’Donnell et al., 2025, 2026), i.e., regions larger than 30 kb showing a presence/absence polymorphism and containing a Captain plus at least one *Starship*-related gene (Fig. 3, bottom panel). We then mapped reads of all other genomes, sequenced with short reads, on these *Starship-*like regions, and used the mean coverage as an indicator of individual *Starship* presence (Fig. 3).

The clonal lineages commercially used for cheese making had a higher content of *Starship*-like regions than the more natural populations, reaching 27.2% of the genome in *P. camemberti* var. *camemberti,* 29.1% in *P. camemberti* var. *caseifulvum* and 27% in *P. biforme cheesy*, against only 24.6% in *P. biforme natural* and 11% in *P. fuscoglaucum* (Fig. 3). The wild species *P. fuscoglaucum,* which does not occur on cheese, thus showed the lowest *Starship* content. Furthermore, *Starship-*like regions were not shared between all strains from a given genetic cluster, indicating that these elements could be acquired or lost after cluster divergence. The highest variability was found in the wild populations *P. biforme natural* and *P. fuscoglaucum,* but striking variations were also observed in the clonal clusters *P. camemberti* and *P. biforme cheesy.* The *P. biforme natural* population showed the greatest variation in terms of *Starship* content, with some strains carrying twice as much *Starship-*like material than others (Fig. 3, right panel). Shared *Starships* were not always located in homologous regions between and within clusters, indicating their acquisition by independent horizontal transfer events or their movement after acquisition.

#### *Rattus*: a *Starship* largely restricted to the *P. biforme cheesy* population and nested within a horizontally transferred *Starship*

A 20kb *Starship* was nearly specific to the *P. biforme cheesy* population, although present in a few *P. biforme natural* strains (Fig. 3). This *Starship* was nested within a much larger *Starship,* spanning 160kb and shared between *P. biforme, P. camemberti* and *P. salamii* (Fig. 4A-B). We named the 160-kb *Starship Bilge* because it carries the cheese-specific *Starship Rattus*, named for its cargo of putative antagonism-related genes (see below) and its mobilization by *Bilge*. *Rattus* contained ten genes, five of which received no functional annotations using Interproscan, Funannotate or Eggnog-scanner. Putative functions were subsequently assigned to three of these genes based on protein structural similarity analyses (Fig. 4C-E; FigSX.protein_comps_full_all). The seven genes with putative functions included the captain (DUF3435), a GH18 secreted endochitinase, a hemolysin-III-like protein, a hemolysin-DF-like protein, a RasGEF domain containing protein, two transcription factors and an osmotin-like protein (allergen *Aspergillus* f4) (Fig. 4B-E; FigSX.protein_comps_full_all). The putative boundaries of *Rattus*, defined by manually identifying direct and terminal inverted repeats at both ends, indicate an unusual *Starship* architecture in which a protein-coding gene (a fungal-specific transcription factor; IPR021842/ENOG503P5WH) lies closer to the element edge than the Captain gene. This contrasts with the canonical *Starship* structure, where the Captain gene is the first coding sequence adjacent to the element boundary. Evidence that this fungal-specific transcription factor constitutes part of the *Rattus* cargo includes its complete absence from long-read assemblies in which *Rattus* is absent (*P. camemberti* var*. camemberti*, *P. fuscoglaucum*, and *P. camemberti* var*. caseifulvum*).

**Fig. 4.**
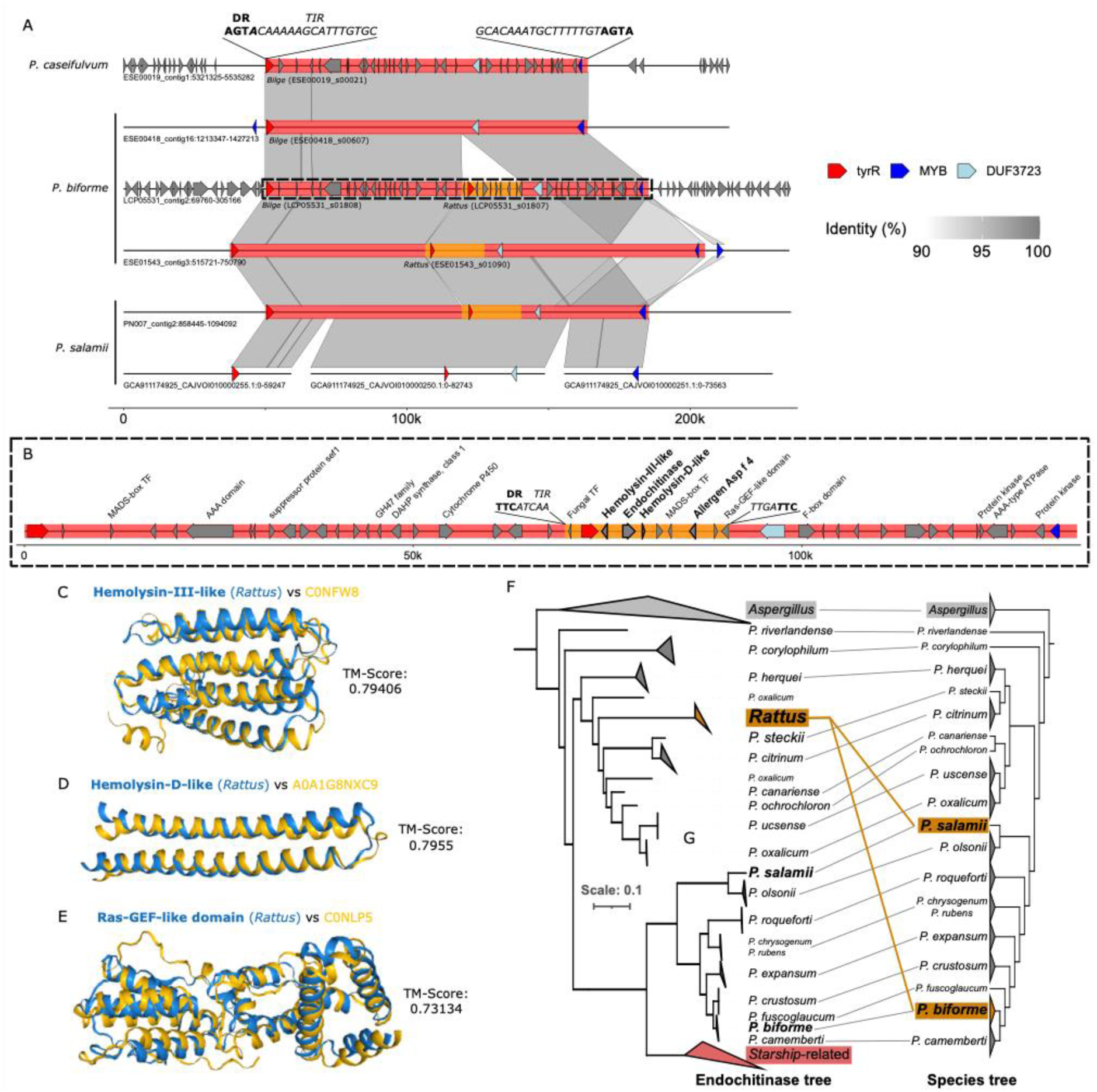
*Rattus:* a *Starship* nearly specific to the *Penicillium biforme cheesy* population, nested within the *Bilge Starship.* (A) The *Bilge Starship* is identical in sequence in *P. camemberti, P. biforme* and *P. salami*, but is located in non-homologous regions, as it was horizontally transferred. Each line represents the scaffold carrying the *Starship* in different strains and species sequenced in long-reads, namely *P. camemberti* var. *caseifulvum* ESE00019*, P. biforme natural* ESE00418 and LCP05531, *P. biforme cheesy* ESE01543 and *P. salami* PN007 and CBS135397 (GCA911174925). Terminal inverted repeats (TIR) and direct repeats (DR) are flanking *Bilge* (motifs indicated at the top of the figure)*. Rattus* is highlighted in orange and is only present in *P. biforme cheesy* and *P. salami*; *Bilge* is highlighted in red and present in all strains presented in this figure. The three genes specific to *Starship-like* regions are indicated with arrows for each strain/species: red for the tyrosine recombinase (*Captain,* DUF3435), dark blue for MYB, and light blue for DUF3723. Genes with putative functions are indicated in the dashed rectangle at the bottom of the figure; the functions of other genes are unknown. The dotted black box highlights the *Bildge-Rattus* region enlarged in (B). Functional annotations given to genes within *Blidge* and *Rattus* are labelled above gene models. Putatively antagonistic genes in *Rattus* are in bold. DR and TIR sequences are shown for *Rattus* as for *Bilge.* (C-E) Alphafold3 predicted structures of three *Rattus* proteins with structural similarities to Foldseek databases. Only alignable portions of each protein structure are shown. (F) On the left is a maximum-likelihood phylogenetic tree placing the *Rattus* endochitinase gene among other endochitinase genes (called core genes). Nodes were collapsed if they were a core endochitinase from the same species, complex or *Aspergillus*. Nodes were also collapsed for the three *Rattus* endochitinase copies, and all other endochitinase proteins found within *Starships* or SLR (‘Starship-related’). On the right is a neighbor-joining k-mer-based species tree of genomes from the same protein datasets used for the endochitinase tree. Nodes are collapsed for multiple genomes of the same species. Lines connect species placement in the phylogenomic tree at right to endochitinases from the same species at left, including core (grey) and *Rattus* (orange) endochitinases.

In the long-read reference assembly of the ESE01543 *cheesy* strain, *Rattus* was flanked by two additional *Starship-*like regions, of 256kb and 69kb, respectively (cluster16 and cluster10 *Starship-*like regions on Fig. 3). These flanking regions are part of another *Starship* element, named *Bilge,* which occurs more broadly than *Rattus* itself, including in all *P. camemberti* strains (both varieties) and the ESE00418 *P. biforme natural* strain (Fig. 4A). *Bilge* was found in non-homolous regions between strains and species, indicative of horizontal transfer, or at least movements within genomes. Notably, the entire large element including *Bilge* and *Rattus* nested within it, nearly restricted in our dataset to *P. biforme cheesy,* was also found in *P. salamii* with >99.5% identity. This suggests a recent horizontal transfer between these species, with *Rattus* having hitchhiked within *Bilge.* Neither *Bilge* or *Rattus* were identified in the previous study having reported *Starships* shared between dry-cured meat *Penicillium* fungi, some being also found in *P. biforme* (Lo et al., 2023).

In the ESE00418 *P. biforme natural* strain, *Bilge* appears to have been inserted adjacent to a large and complex *Starship-*like region (SLR24; Fig. 3). This region contains a Captain gene associated with a MYB-like DNA-binding protein-coding gene, a genomic organization commonly observed in *Starships*. The region likely represents a composite structure formed through multiple nesting events as it contains four captains, four MYB-like genes and four DUF3723 domains. This *Starship-*like region spans approximately 1.5 Mb and contains 1,979 genes (Fig. 3: SLR24). The region was found entirely (coverage percentage >99%) in only two *natural P. biforme* strains (ESE00418, ESE01960) and three *cheesy P. biforme* strains (ESE00087, ESE00089, ESE01262). It was never found complete in *P. camemberti* (mean coverage percentage of *P. camemberti* strains: 57.9%) or *P. fuscoglaucum* (mean coverage percentage of *P. fuscoglaucum* strains: 28.3%).

The GH18 secreted endochitinase predicted in *Rattus* may contribute to adaptation to cheese-making environments by enhancing competitive interactions with other fungi colonizing cheese rinds. By degrading fungal cell walls, it could help exclude competitors and promote rapid colonization of the cheese surface. To investigate the origin of the *Rattus*-encoded endochitinase, we reconstructed a protein tree including all GH18 endochitinase-encoding proteins identified in our long-read assemblies and a set of publicly available *Penicillium* and *Aspergillus* genomes. From our long-read assemblies, endochitinase proteins located within *Starships* or SLRs were designated as *Starship*-related proteins. The tree shows a set of conserved core secreted endochitinase B1 proteins that follows the species tree (Fig 4F; FigSX.endochitinase.phylogeny.full.mod). In contrast, the *Rattus* endochitinase branches are distant from the core endochitinase from *P. biforme* and *P. salami*, further supporting its origin as a horizontal gene transfer rather than as a gene duplication. All other *Starship*-related endochitinases formed a monophyletic clade also discordant with the species tree, and distant from the *Rattus* endochitinase. The tree also shows that, among three copies of these endochitinases present in the reference genome of *P. oxalicum* (GCA_001723175.3), a single one branches as in the species tree, the two other copies being distant, also suggesting a *Starship* origin rather than a gene duplication. A similar approach was applied to the Allergen Asp F 4 protein also present in *Rattus*; the placement of the *Rattus* gene in discordance with the species tree also suggested a horizontal gene transfer (FigSX.AspF4_tree.asp_rooted.mod). The other cargo genes in *Rattus* contained no detectable homologs in the same genomes, but they all displayed functions also likely involved in antagonistic interactions.

### Phenotypic differentiation between *P. biforme cheesy* and *natural* populations

To test whether the *P. biforme cheesy* population has been domesticated for cheesemaking, we investigated to what extent it evolved specific adaptive traits for cheese. We measured several traits potentially involved in adaptation to cheese or on which selection could be relaxed in this environment, using 10 strains from *P. biforme cheesy*, *P. biforme natural*, and the wild relative *P. fuscoglaucum.* Indeed, fast-growing fungi promote faster cheese maturation and create an attractive rind. They can also help prevent spoilage caused by bacteria, yeasts and molds. This is exemplified by *P. camemberti,* which is capable of rapidly covering the surface of soft cheeses. Here, we grew strains on various media at different temperatures and measured the growth area every 7 days until 21 days. We used a cheese medium, a minimal medium to mimic the harsh conditions found in wild environments, and malt extract agar to determine whether strains metabolize sugars other than those present in milk. Experiments were conducted at two temperatures: 12°C to simulate cheese cave conditions, and 25°C, the optimal growth temperature for *Penicillium* species. We found that *P. biforme cheesy* generally grew faster under conditions mimicking cheese ripening than the wild *P. fuscoglaucum* species (Fig. 5A), as expected following domestication. Indeed, Kruskall-Wallis tests indicated that *P. biforme cheesy* grew more rapidly than *P. biforme natural* or *P. fuscoglaucum* at 25°C (Fig. 5A). In cave conditions, at 12 °C in the dark, *P. biforme cheesy* grew faster than *P. fuscoglaucum* on cheese (Wilcoxon test, p=0.002, Table S2), with no significant difference between *natural* and *cheesy* populations. At 25°C, *P. biforme cheesy* grew faster than *P. biforme natural* (Wilcoxon test, p=0.008, Table S2) and *P. fuscoglaucum* (Wilcoxon test, p=0.001, Table S2). However, the full model in the ANOVA only showed a significant effect on growth rates of the culture medium, the temperature and their interaction (Table S2), and no significant overall effect on growth of the species/population.

**Fig. 5.**
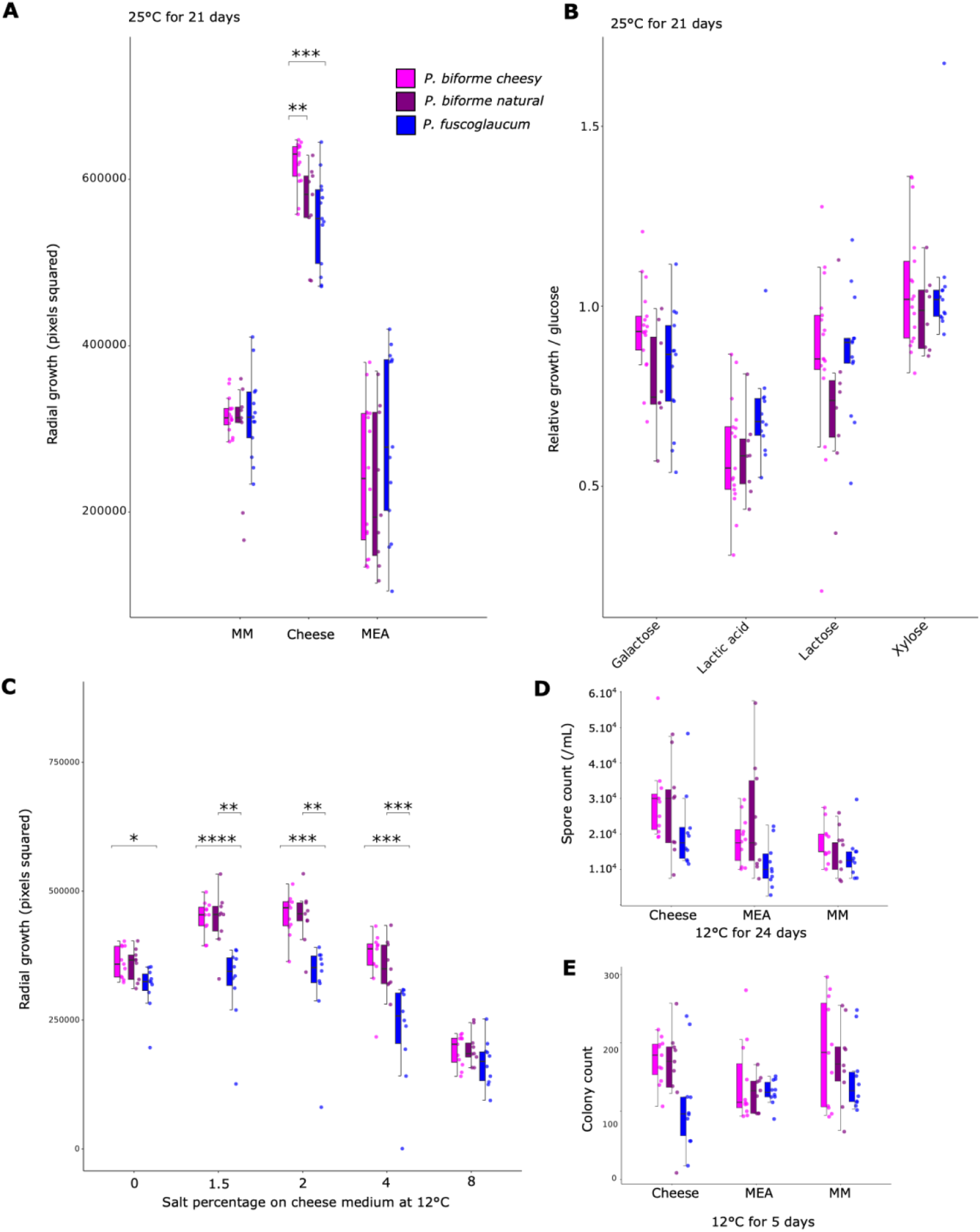
Measures of different phenotypes of *Penicillium biforme cheesy* and *natural* populations, as well as of *P. fuscoglaucum,* in various conditions. A) Radial growth at 25°C on minimal medium (MM), cheese (1.5% salt) and malt extract agar (MEA) media for 21 days; B) Relative growth on various carbon sources (normalized to growth glucose): galactose, lactic acid, lactose and xylose. Measurements were done after 21 days of incubation at 25°C ; C) Radial growth on a cheese medium with different salt concentrations, from 0 to 8% ; D) Number of spores on minimal medium (MM), cheese and malt extract agar (MEA) media after 24 days of growth at 12°C ; E) Colony count after 5 days of growth at 12°C on minimal medium (MM), cheese and malt extract agar (MEA) media. Each dot represents a strain. Only significant differences between populations/species are shown (Wilcoxon tests; * = p <= 0.05; ** = p <= 0.01; *** = p <= 0.001; Table S2). The color indicates assignments to the three populations/species as in other figures: pink, *P. biforme cheesy ;* purple, *P. biforme natural* ; blue, *P. fuscoglaucum*.

Our experiments further indicated that *P. biforme cheesy* and *natural* displayed enhanced tolerance to salt (Fig. 5C), consistent with adaptation during domestication for cheese production. Indeed, high concentrations of salt are used in cheese to avoid food spoiler micro-organisms. At any salt concentration up to 8%, being twice the maximum percentage of the saltiest cheese (Roquefort), all *P. biforme* populations grew faster than *P. fuscoglaucum* (Fig. 5C, Table S2). This effect was even stronger at 1.5, 2 and 4% of salt, which corresponds to the actual range of salt concentration of cheese.

We also found that cheese-associated *P. biforme* strains exhibited carbon source usage consistent with adaptation to the cheesemaking environment. On galactose and lactose, the *P. biforme cheesy* population tended to grow faster than *P. biforme natural*, which could reflect adaptation to cheese sugars released from lactose hydrolysis by bacteria early in ripening, but the difference was not significant. *Penicillium biforme cheesy* grew significantly less than its wild relative *P. fuscoglaucum* on lactic acid (Wilcoxon test, p=0.004), which is abundant early in ripening, but largely depleted by the time *Penicillium* molds begin to grow. There was no difference between the two *P. biforme* populations in terms of growth on lactic acid (Fig. 5B; Table S2). Growth on xylose, a plant sugar, did not differ between populations (Fig. 5B; Table S2).

*Penicillium biforme cheesy* and *natural* on cheese medium produced significantly more spores than the wild *P. fuscoglaucum* (Fig. 5D, Table S2). Cheese strains produced higher amounts of spores than wild strains, which is beneficial for obtaining spores for cheese inoculation, and for efficiently colonizing the cheese surface, thereby inhibiting spoiler microorganisms. Colony counts after five days, used as a proxy for germination rate, were also higher in the *cheesy* population than in *P. fuscoglaucum* (Fig. 5E, post-hoc Tukey test, p = 0.04), with no difference between *cheesy* and *natural P. biforme* (post-hoc Tukey test, p = 0.7) (Fig. 5E, Table S2). An ANOVA confirmed significant effects of species and culture medium on both spore production and colony count, as well as of strain origin on spore production (Table S2).

Mold fluffiness contributes to cheese surface coverage and texture, and has likely been selected during domestication. However, cheese-associated *P. biforme* did not differ in fluffiness from *natural P. biforme* or *P. fuscoglaucum* (Table S2). Only *P. camemberti* was clearly fluffier than all other populations, as previously reported, with var. *caseifulvum* less fluffy than var. *camemberti*, but still fluffier than *P. biforme* and *P. fuscoglaucum* (Fig. S1).

By monitoring lipolytic and proteolytic activities weekly, we found that *P. biforme cheesy* showed higher lipolytic activity at 28 days than the *natural* population (Fig. 6B, Wilcoxon signed-rank test, *p* = 0.02) and no significant difference with *P. fuscoglaucum* was found. This result is consistent with a *P. biforme cheesy* adaptation to cheese, where fat breakdown contributes to flavor and aroma development. For proteolytic activity, which affects both texture and flavor by breaking down proteins, no difference was observed between *cheesy* and *natural P. biforme* during the first 21 days, but proteolytic activity tended to be lower at day 28, while the wild *P. fuscoglaucum* showed no proteolytic activity (Fig. 6A, Table S2) during 28 days.

**Fig. 6.**
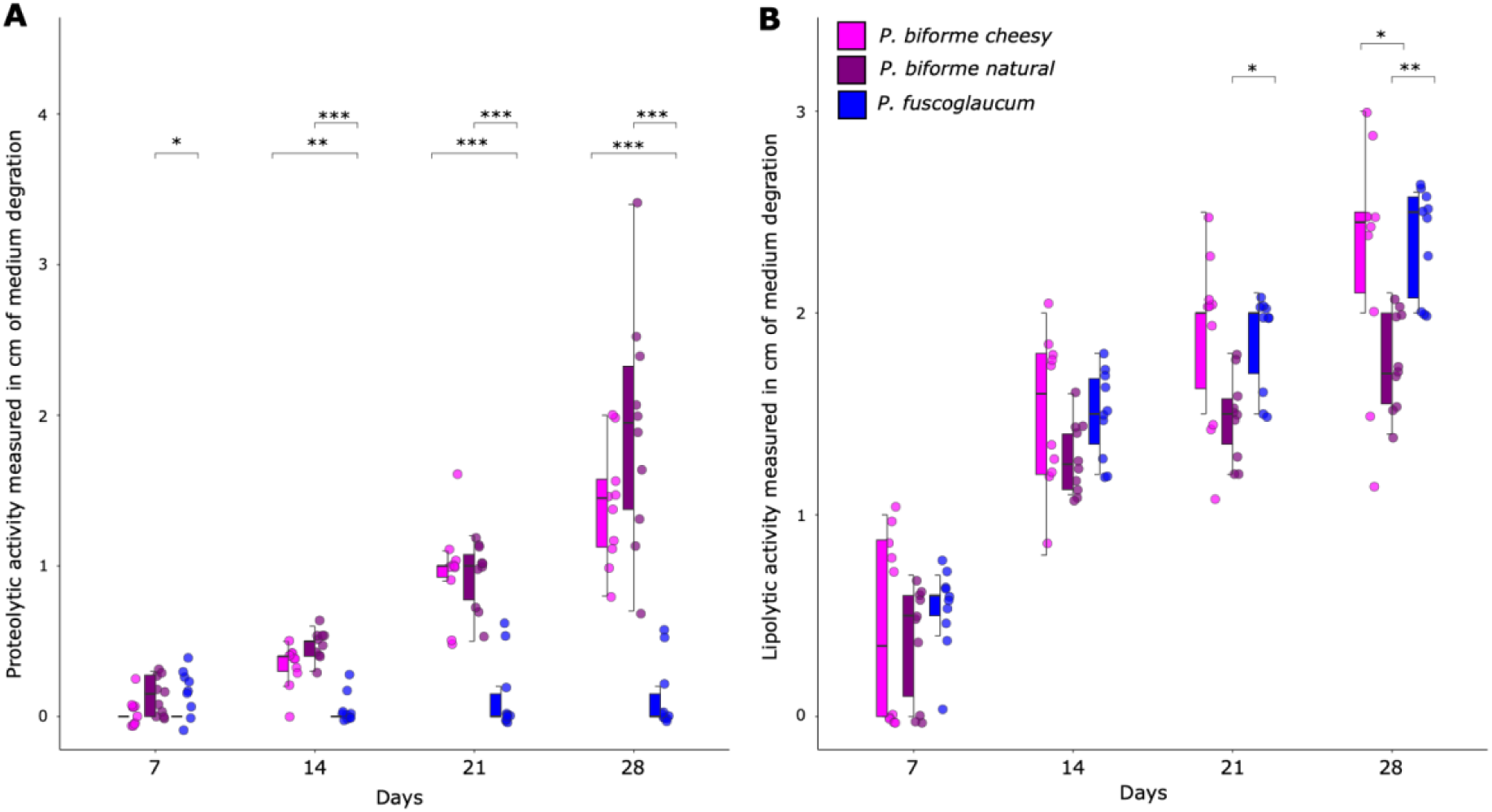
Lipolytic and proteolytic activities of *Penicillium biforme* and *P. fuscoglaucum,* estimated by weekly monitoring cumulated medium degradation for 28 days at 12°C. A) Proteolytic activity measured on agar semi-skimmed milk; B) Lipolytic activity measured on agar tributyrin. Each dot represents a strain, and colors indicate the cluster of origin: pink for *cheesy*, purple for *natural* and blue for *P. fuscoglaucum.* Only significant comparisons are shown (Wilcoxon tests; * = p <= 0.05; ** = p <= 0.01; *** = p <= 0.001; Table S2).

Mycotoxins are fungal metabolites that contribute to ecological interactions, for example by inhibiting competing microorganisms but can also represent undesirable substances in food products. In domesticated molds, the ability to produce mycotoxin can be lost by positive or relaxed selection, or it could be increased if this is beneficial for excluding undesired contaminating molds. Both *natural* and *cheesy P. biforme* populations produced cyclopiazonic acid (CPA) on YES medium, with significantly higher production in the *cheesy* than in the *natural* population (Wilcoxon rank-sum test, p = 0.009; Fig. S2). In contrast, *P. camemberti* var. *caseifulvum* cannot produce CPA, which we previously showed was due to a frameshift in the biosynthesis cluster (Ropars et al., 2020), whereas *P. camemberti* var. *camemberti* is a known CPA-producer (Le Bars, 1979; Ropars et al., 2020). Cheese is however less conducive to toxin production than YES medium.

By inhibiting competitor microorganisms, *P. biforme* can help exclude food spoilers in cheese. By inoculating challengers on a *P. biforme* lawn, we showed that it effectively inhibited the growth of *Mucor racemosus, P. fuscoglaucum, Scopulariopsis asperula,* and *Yarrowia lipolytica*, with the *cheesy* population showing stronger inhibition ability than the *natural* population (post-hoc Tukey, p = 0.04; Fig. 7, Table S2, the ANOVA showing significant effects on challenger growth of species challenger, species lawn and of their interaction). No inhibition occurred when challengers were physically separated by using splitted Petri dishes, indicating that the effect does not rely on volatile compounds (Table S2).

**Fig. 7.**
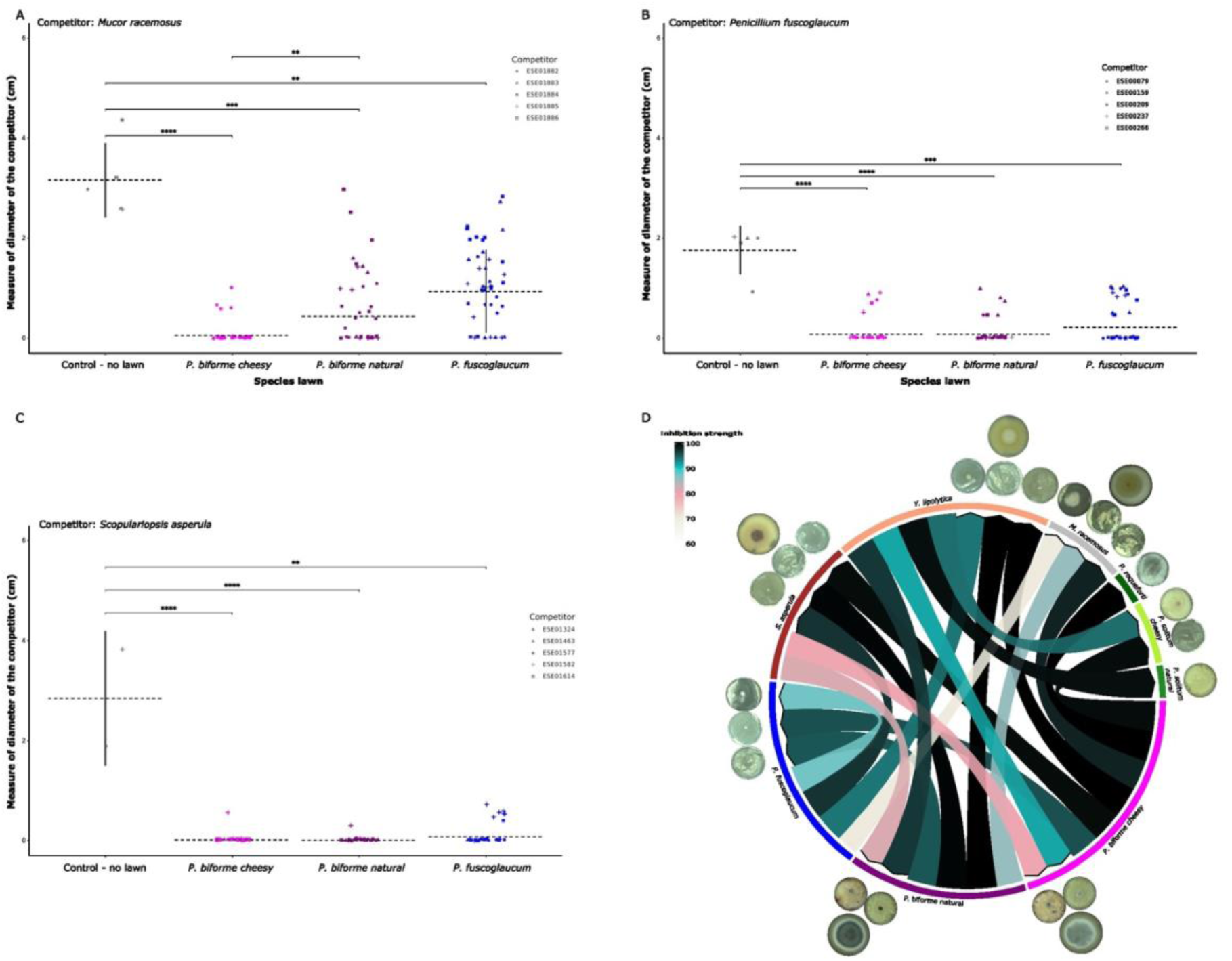
Growth inhibition experiment. A-C) Competition experiments with *Penicillium biforme* and *P. fuscoglaucum* strains as lawns, organized by species/clusters (x-axis); Competitors were inoculated 24 hours after the lawn as a central point on cheese medium: *Mucor racemosus* (panel A)*, Penicillium fuscoglaucum* (panel B, only strains with white colonies were selected to have a contrast between the lawn and the competitor) and *Scopulariopsis aperula* (panel C). Control Petri dishes (black dots) only had competitors, i.e. without lawns. The dot shape corresponds to the competitor strain. Only significant differences between populations/species are shown (Wilcoxon tests; * = p <= 0.05; ** = p <= 0.01; *** = p <= 0.001; Table S2). D) A summary of all interaction tests represented in a Circos plot, with arrows indicating the direction of inhibition: the tail represents the species used for the lawn, and the arrowhead represents the competitor species. The color indicates the inhibition strength estimated as the mean growth calculated per species challenger, normalized by the control species challenger (Petri dish without species lawns) and transformed in percentage.

To assess overall phenotypic differentiation between populations and species, we performed a principal component analysis integrating eleven phenotypic traits measured across the 30 strains belonging to *P. biforme cheesy, P. biforme natural* and *P. fuscoglaucum*, including growth, salt tolerance, carbon utilization, sporulation, germination, lipolytic and proteolytic activities. *Penicillium biforme* appeared clearly distinct from the wild *P. fuscoglaucum* species, and the *cheesy* and *natural P. biforme* also appeared to have different phenotype ranges, although to a lesser extent. *Penicillium fuscoglaucum* was clearly separated from all *P. biforme* along the first principal component, while the *cheesy* population displayed a slight shift relative to the *natural* one along the second component (Fig. 8). The first and second components were positively associated with growth on cheese at all tested salt concentrations (1.5–8% NaCl) and temperatures (12°C and 25°C), indicating selection for faster growth on cheese in *P. biforme*, and perhaps more so in the *cheesy* population. The first component, separating *P. fuscoglaucum* from *P. biforme*, was also positively associated with spore production, germination rate, and proteolytic activity, all of which being key traits for cheesemaking.

**Fig. 8.**
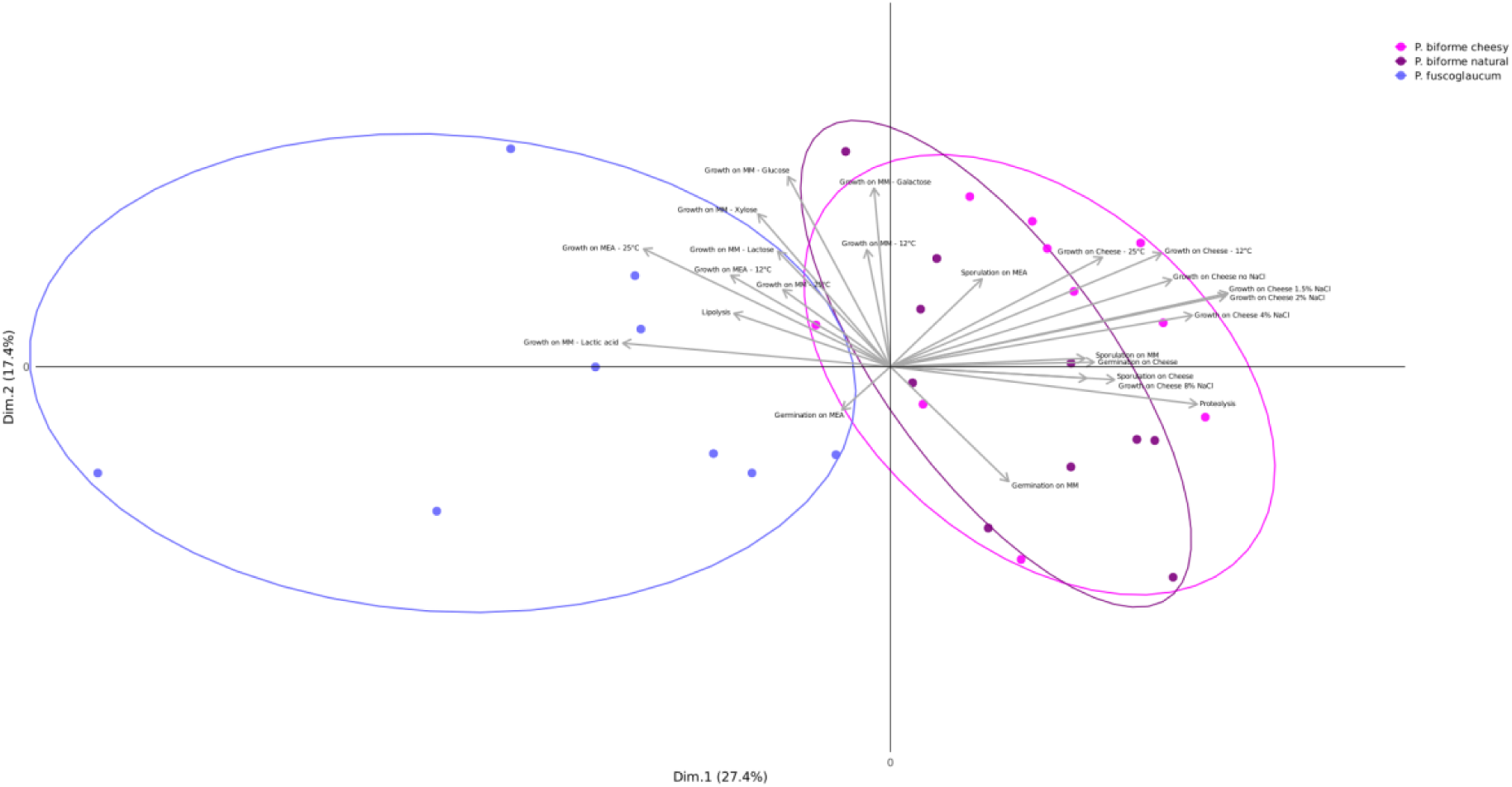
Principal component analysis (PCA) illustrating the phenotypic differences between *Penicillium biforme* populations and *P. fuscoglaucum.* The association between the two first PCA axes and the variables are shown. Variables: growth at 12°C and 25°C on cheese, minimal medium (MM) and malt extract agar (MEA), growth on cheese at different salt concentrations (0, 1.5, 2, 4, 8% NaCl), growth on MM supplemented with different carbon sources (glucose, xylose, lactose, lactic acid, galactose), proteolysis, lipolysis, spore production (sporulation) on MM, MEA and cheese, germination rate on MM, MEA and cheese.

### Volatile organic compounds reveal domestication-associated aroma signatures

Volatile organic compounds (VOCs), which substantially contribute to cheese flavor development and are primarily generated through lipolytic and proteolytic activities (Bertuzzi et al., 2018; Molimard & Spinnler, 1996), differed significantly among populations (PERMANOVA, *df* = 4, *p* = 0.006). Pairwise comparisons indicated that VOC profiles of *P. biforme cheesy* differed significantly from those of the wild species *P. fuscoglaucum*, whereas a substantial overlap was observed between *cheesy* and *natural* populations (Fig. SX_volatiles_NMDS Fig. SX_Volatiles_heatmap_pvalues_adonis). VOC composition showed considerable within-population variability, particularly in *P. biforme natural*, but also within the clonal domesticated populations, i.e., *P. biforme* cheesy and the two varieties of *P. camemberti,* var. *camemberti* and var. *caseifulvum* (Fig. 9A; Fig. SX_alphaDiv; Fig. SX_volatiles_NMDS).

**Fig. 9.**
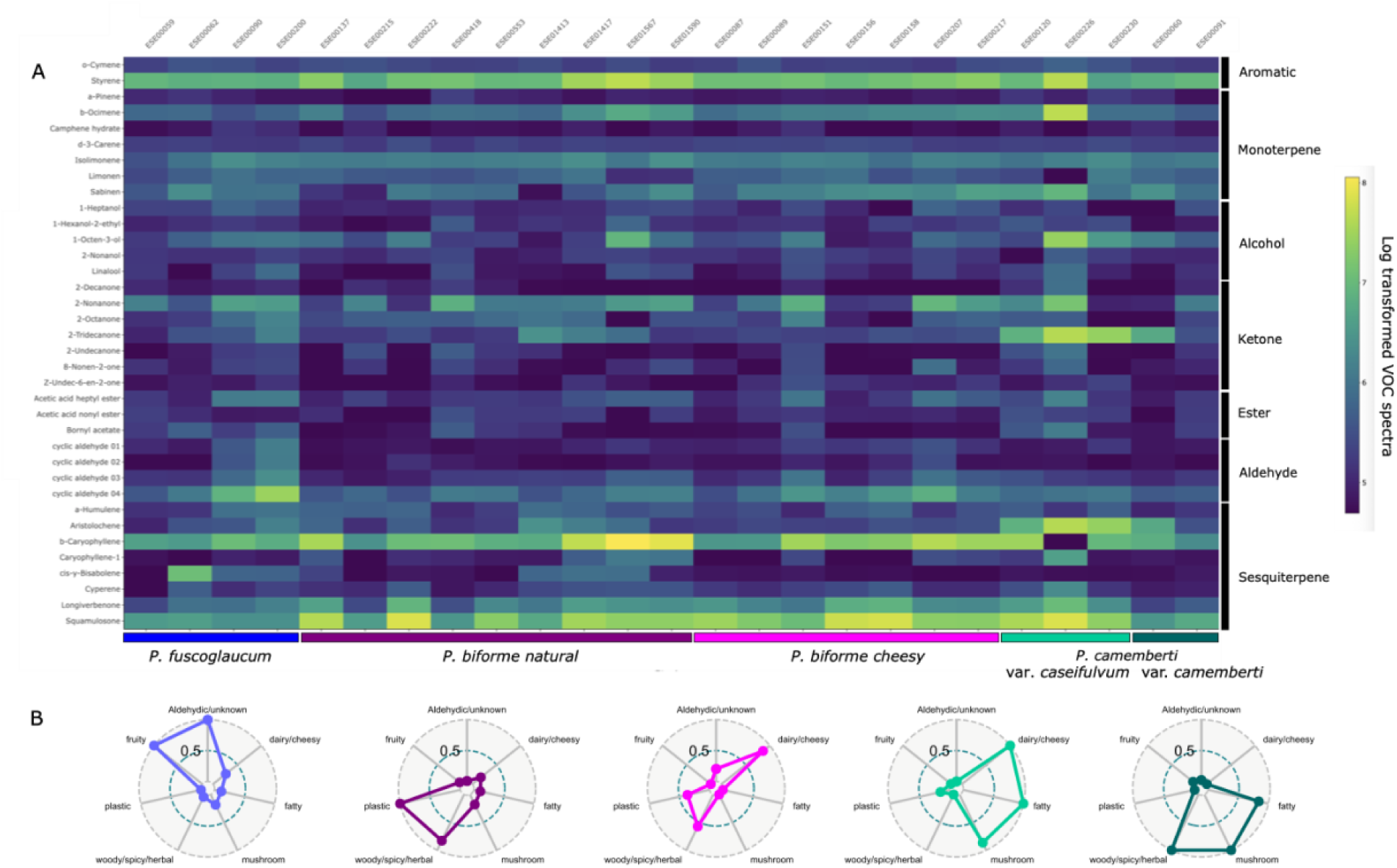
Volatile organic compounds (VOCs) and aroma-family composition of VOCs produced by *Penicillium biforme, P. camemberti* and *P. fuscoglaucum* grown on a cheese medium for 21 days. A) Heatmap representing the log-transformed values of volatile emissions. Volatile organic compounds (VOCs) are shown as peak area per gram fresh weight and are classified according to their chemical classes, including aromatic compounds, monoterpenes, alcohols, ketones, esters, aldehydes and sesquiterpenes, as indicated at right. B) Radar plots representing the aroma-family composition of volatile organic compounds. Seven aroma families were defined using public databases, including three relevant for cheese-making (dairy/cheesy, fatty, mushroom); VOC abundances were expressed as proportions of the total VOC signal, averaged across replicates, and then further averaged across populations. The contribution to each aroma family was taken as the sum of the VOC abundance for the focal family. For visualisation purposes, the values for each aroma family were normalized across populations to a 0–1 scale.

Random Forest classification revealed moderate discrimination between species (out-of-bag error = 60%), although terpenoid compounds, particularly sesquiterpenes, strongly contributed to population differentiation (Table SX MeanDecreaseAccuracy and MeanDecreaseGini; Fig. SX ROC-AUC analyses; Table SX ROC-AUC analyses). OPLS-DA integrating VOCs and genomic data likewise identified terpene metabolism as a major source of species differentiation (Fig. SX_table_OPLS_summary). VOC differentiation was not significantly correlated with genetic distance (Mantel test: r = 0.226, p = 0.085), indicating that factors beyond shared ancestry substantially contribute to volatile variation.

Several volatile compounds were enriched in *P. biforme* cheesy or *P. camemberti* var. *caseifulvum* and were absent or strongly reduced in the wild *P. fuscoglaucum*: squamulosone and longiverbenone (all *P. biforme* and *P. camemberti* var. *caseifulvum*), β-caryophyllene (both *P. biforme* clusters), 2-tridecanone and aristolochene (both *P. camemberti* varieties), d-3-carene (*P. camemberti* var. *caseifulvum* and *P. biforme* cheesy), styrene (var. *caseifulvum* and *P. biforme*), and 8-nonen-2-one (var. *caseifulvum*). These differences suggest metabolic shifts associated with adaptation to cheese environments.

To facilitate biological interpretation of VOC production, we performed an analysis per aroma family, after excluding styrene (as usually done because it is a well-documented plastic laboratory equipment contaminant) and the four unidentified cyclic aldehyde features. VOCs were assigned to seven aroma sensory families using two public databases (https://www.thegoodscentscompany.com/ and https://scent.vn/en). Aroma family profiling revealed clear differences between lineages (Fig. 9B). In particular, the two varieties of *P. camemberti*, as well as *P. biforme cheesy,* showed greater contributions of cheese-like aroma families, i.e. dairy/cheesy, fatty and mushroom, whereas *P. fuscoglaucum* and *P. biforme natural* displayed reduced representation of these cheese-associated signatures (Fig. 9B).

## Discussion

*Penicillium biforme* is recurrently found in the cheese environment but its wild, ancestral-like ecological niche is still unknown. In a previous study, we showed that the emblematic white mold *P. camemberti* has been domesticated, as this lineage has undergone a drastic reduction in genetic diversity, as a result of very strong selection for traits beneficial for cheese making, including white color and fluffy morphology (Ropars et al., 2020). Previous studies also suggested that *P. biforme* has adapted to cheese and dry-cured meat production, as we could identify specific traits that have evolved for food adaptation when compared to its wild relatives (Lo et al., 2023; Ropars et al., 2020). At that time, however, we only had 25 *P. biforme* genomes, mostly isolated from cheese. Here, we substantially expanded our sampling by including new *P. biforme* strains isolated from artisanal or industrial cheeses, as well as non-food environments, enabling a broader ecological perspective on the evolutionary history of *P. bifor*me.

### Genomic analyses reveal a new domesticated, clonal lineage in *P. biforme*

By using a total of 75 *P. biforme,* 22 *P. camemberti* and 22 *P. fuscoglaucum* genomes, we showed that *P. biforme* was divided into three genetic clusters: *P. camemberti, P. biforme cheesy* and *P. biforme natural.* We thus identified a new *P. biforme* lineage, distinct from *P. camemberti,* which we named *cheesy. Penicillium biforme cheesy* displayed footprints of selection for cheese making: *cheesy* only contained strains of a single mating type, had very low genetic diversity (as low as the clonal lineage *P. camemberti*), carried high numbers of *Starship* elements and displayed specific traits beneficial for cheese production, compared to other *P. biforme* strains and to its wild relative *P. fuscoglaucum.* These traits included faster growth on cheese, galactose and lactose media, higher sporulation and germination rates on cheese, higher lipolytic activity, better inhibition capacities and specific VOCs. Strains belonging to the *P. biforme cheesy* lineage were collected from non-artisanal cheeses or dried sausages, suggesting that they have been inoculated as commercial cultures. *Cheesy* is thus likely a clonal lineage cultivated and sold in the industry, for making cheese and dry-cured meats. Some fungal spore producers do sell commercial mold cultures as *P. biforme*.

In contrast to the *cheesy* population, the genetic diversity in *P. biforme natural* was high, with no genetic differentiation between cheese-associated strains and those isolated from other environments, such as hay or lacto-fermented cucumber. Additionally, we isolated genetically distinct strains from artisanal cheeses produced by the same producers. Taken together, these findings indicate that the natural *P. biforme* population is wild; it may come from a plant-related environment, as *P. biforme* strains isolated outside food environments are typically associated with plants. .

Both *P. camemberti* and *cheesy P. biforme* are nested within *P. biforme*, which is therefore not monophyletic. However, *P. camemberti* being a commercially important culture in the cheese industry, the two taxonomic names *P. camemberti* and *P. biforme* are often retained, even though they actually correspond to the same species.

The evolutionary history of *P. biforme* looks very similar to that of *Geotrichum candidum,* also naturally colonizing a variety of cheeses and displaying high genetic diversity, with the exception of a few clonal lineages being inoculated in cheeses (Bennetot et al., 2023). We identified strong and recent signatures of selection for cheesemaking in both species, in the cheese_2 lineage in *G. candidum* (Bennetot et al., 2023) and in the *P. camemberti* and *cheesy* lineages in *P. biforme* ((Ropars et al., 2020) and this study). In both species, the cultivated lineages, widely used in cheesemaking and under strong selection, exhibit a severe loss of genetic diversity, as previously also found in the blue-cheese mold *P. roqueforti* (Dumas et al., 2020).

### *Starship* mobile elements are abundant and generate variability

Single nucleotide polymorphism is not the only component of genetic variation in populations. In fungi, horizontal gene transfers are recognized as important drivers of adaptation. Notably, a superfamily of giant transposable elements has recently been identified in filamentous fungi, named *Starships,* exhibiting high mobility between genomes and even across species, and able to move dozens of cargo genes (Urquhart et al., 2024). Here, our high-quality genome assemblies allowed accurately identifying *Starships,* revealing that they were an important source of variability, even within populations with little SNP diversity. Indeed, we could identify variation in terms of *Starships* presence/absence in *P. biforme*, especially in the three domesticated lineages, despite clonality. *Starships* have in fact been recognized as drivers of variability and adaptation in several clonal fungal pathogens, such as in the human pathogen *Aspergillus fumigatus* (Gluck-Thaler et al., 2025), in the plant pathogens *Verticillium dahliae* (Sato et al., 2025) and *Fusarium xylarioides* (Peck et al., 2024), with *Starship-*mobilized genes encoding functions important for virulence. *Starship-*like regions have accumulated in clonal, domesticated lineages of *Penicillium* and *Aspergillus* species (O’Donnell et al., 2025), further suggesting a role in rapid adaptation in non-recombining populations. Here, we identified more *Starship-like* regions in the recombinant *P. biforme natural* than in its wild relative *P. fuscoglaucum.* This new finding further supports the view that *Starships* play a key role in shifting lifestyles and in promoting genetic exchange, and can do so even in wild populations.

Focusing on *P. biforme,* we detected presence/absence *Starship* polymorphism within domesticated populations. Most importantly, *P. biforme* had a higher *Starship* content than its wild relative *P. fuscoglaucum*, in agreement with the increase in *Starship* content of domesticated fungal lineages that we previously detected (O’Donnell et al. 2025). Notably, we identified a small *Starship* specific to the *P. biforme cheesy* population of 20kb, containing 10 genes with some putative functions likely in antagonistic interactions, i.e. GH18 endochitinase, hemolysin-III-like, hemolysin-D-like and an osmotin-like protein (allergen *Asp* f4). This *Starship* was nested within a much bigger *Starship* of more than 100kb, and we called them *Rattus* and *Bilge*, respectively. In LCP05531, we identified, adjacent to *Bilge,* the largest *Starship*-like region ever described, likely made up of four juxtaposed individual *Starships*, encompassing and 1,773 genes across 1,409,000 bp in total, with several genes encoding protein kinases, iron transport, lipid transport and metabolism, lactose/galactose, carbohydrate metabolism and transport. This region was present in all *P. biforme* strains, although with various degrees of completeness (39-100%), being found in full length in three *cheesy* strains and two *natural* ones. This *Starship*-like region was present in all *P. camemberti,* with various but relatively high degrees of completeness (43-70%), whereas it was absent or only at low levels of completeness (35% maximum) in the wild relative *P. fuscoglaucum*. Altogether, this provides compelling evidence for the role of this particular *Starships* in cheese adaptation. Here, we thus add novel evidence of the important role of *Starships* in rapid adaptation to made-made environments.

By comparing the endochitinase and Allergen Asp F 4 cargo genes from *Rattus* with core homologs, we have provided further evidence for a horizontal transfer of these *Starship*-like regions and perhaps some indications on their origin. The other cargo genes in *Rattus* had no homologs in the same genomes; strikingly, they all appear to display predicted functions involved in antagonistic interactions. Taken together, these findings suggest that *Rattus* is a horizontally transferred adaptive module that has moved across distantly related species, as supported by the phylogenetic placement of its endochitinase and Asp F4-like genes. Its maintenance and spread may be driven by the overall competitive advantage conferred by its cargo gene repertoire.

The *Rattus* cargo gene with the most prominent role in antagonism is the endochitinase B1, and the gene tree shows that this gene copy has been acquired from a distant species. Within the same *Penicillium* clade, *P. oxalicum* also contained two additional copies of this endochitinase, two likely acquired by HGT, and one copy being sister to the *Rattus* endochitinase. In *P. oxalicum*, the endochitinase B1 has been shown to contribute to antagonistic interactions against other fungi (Xie et al., 2021) and insect pathogenesis (Song et al., 2024). The acquisition of a divergent endochitinase by cheese fungi via *Rattus* may enhance competitive ability in cheese-making environments, where rapid exclusion of competitors is likely beneficial.

### The *P. biforme cheesy* lineage has evolved traits beneficial for cheese making

In agreement with the view that the *P. biforme cheesy* population corresponds to a domesticated lineage, we found that it has evolved several traits beneficial for cheesemaking compared with both its wild relative *P. fuscoglaucum* and the *P. biforme natural* population. Indeed, *P. biforme cheesy*, when on cheese medium, grew faster, produced more spores that germinated at a higher rate, likely enabling more rapid colonization of cheese surfaces. Furthermore, *P. biforme cheesy* grew faster on galactose and lactose, two sugars present in milk and cheese. The *cheesy* lineage also exhibited greater lipolytic activity than the *natural P. biforme* lineage. Lipolysis is essential for flavor development because it releases free fatty acids that serve as precursors for several classes of volatile organic compounds (VOCs), including ketones, alcohols and esters, all important contributors to the aroma of surface-ripened cheeses (Bertuzzi et al., 2018; Molimard & Spinnler, 1996).

Our volatilomics analyses indicated that VOC production differed among lineages and contributes to phenotypic differentiation associated with adaptation to cheese. Ketones showed the highest variability within populations and species. Terpenoids, particularly sesquiterpenes, were among the compounds that contributed most strongly to the differentiation between populations. The most significant difference in VOC composition was observed between *P. biforme cheesy* and the wild species *P. fuscoglaucum*, whereas the two *P. biforme* populations still showed substantial overlap. VOC variability was also present within populations, including clonal populations such as *P. biforme* cheesy and both varieties of *P. camemberti*. This pattern is consistent with the broader view emerging from comparative fungal volatilomics: fungal VOC profiles contain both shared and lineage-specific components, they reflect ecological differentiation, and can exhibit high variability within lineages (Guo et al., 2021; Zhu et al., 2026). In our dataset, this combination of partial population differentiation and substantial within-population variability suggests that VOC phenotypes are biologically meaningful and relevant to adaptation to cheese production, and is not determined by genetic relatedness alone. In the blue cheese fungus *P. roqueforti* too, VOC distributions were found differentiated between lineages, and with specific and more diverse VOCs in cheese lineages, even those with little genetic diversity (Caron et al., 2021; Crequer et al., 2024).

More specifically, several VOCs were found in higher concentrations in the domesticated strains *P. biforme* cheesy and *P. camemberti* var. *caseifulvum* but were absent or present in lower concentrations in the wild species *P. fuscoglaucum*. These compounds included squamulosone, longiverbenone, β-caryophyllene, 2-tridecanone, aristolochene, D-3-carene, styrene and 8-nonanone. A cyclic aldehyde was produced by both *P. biforme* cheesy and wild *P. fuscoglaucum*. Of these compounds, 8-nonen-2-one is of particular interest as related methyl ketones and associated compounds have previously been reported in the flavour formation of Brie- and Camembert-type cheeses (Karahadian et al., 1985). More generally, ketones, alcohols, acids, esters, aldehydes, and sulphur-containing volatiles are well-established contributors to the aroma of surface-ripened cheeses (Bertuzzi et al., 2018; Molimard & Spinnler, 1996). Therefore, the clonal *Penicillium* populations inoculated in cheese, namely *P. camemberti* and *P. biforme cheesy*, produce VOCs that likely contribute to cheese flavour and aroma. In addition, by assigning VOCs to aroma families, we found that *P. camemberti* var. *caseifulvum* and *P. biforme cheesy* produced pleasant aromas, typical in cheeses. This was similarly reported in the blue-cheese mold *P. roqueforti,* in which cheeses made with cheese strains produced the characteristic aromas of ripened blue cheeses, while those made with non-cheese strains had unpleasant odors, described as resembling a wet mop (Caron et al., 2021). This supports the view that human selection has shaped fungal traits for cheesemaking, and not only growth and color, but also the metabolic pathways responsible for desirable aroma.

We also found that the *P. biforme* cheesy lineage was more effective at inhibiting other fungal species than the *natural* population, particularly those considered undesirable contaminants by stakeholders, such as *Mucor racemosus*. As no inhibition was observed when the strains were physically separated, this inhibitory capacity is likely to be due to diffusible non-volatile compounds rather than VOC-mediated interactions under the conditions tested here.

While we did not rigorously measure colour in the present study, we observed that the *P. biforme* cheesy lineage was blue-grey, i.e. not completely white like *P. camemberti* var. *camemberti*. Nevertheless, a previous study showed that *P. biforme* was whiter than the wild, closely related *P. palitans* (Lo et al., 2023).

Overall, our findings support the view that the *P. biforme* cheesy lineage has undergone adaptation driven by domestication to the cheese environment, combining enhanced growth and competitive ability with metabolic traits that are likely to contribute to cheese ripening and aroma.

## Conclusion

In conclusion, our study revealed that, in addition to the emblematic white, fluffy mold *P. camemberti* that gives soft cheeses their immaculate appearance, another lineage has been domesticated from *P. biforme* for food fermentation: a clonal, blue-grey lineage that we have named ‘*cheesy*’. This lineage exhibits the classical genomic hallmarks of domestication, including extremely low genetic diversity and a single mating type. It has also evolved a variety of traits advantageous in the cheese environment, including faster growth on cheese and milk-associated sugars, increased sporulation and germination in cheese, stronger lipolytic activity, greater competitiveness against other fungi and distinct volatile profiles that are potentially important for cheese ripening and aroma. Together, these findings support the view that the *P. biforme cheesy* lineage had undergone recent adaptation driven by domestication to cheese and possibly other fermented foods, such as dry-cured meat. Our results also emphasize the role of *Starships* in this process of rapid fungal adaptation. The domesticated lineages differed markedly in *Starship* content, allowing the rapid gain of hundreds of genes potentially involved in metabolism and competition, suggestive of a role in adaptation to cheese. More broadly, our study contributes to the growing body of evidence suggesting that domestication in food-associated fungi can occur rapidly through the combined effect of strong selection on beneficial traits and genome innovation mediated by horizontally transferred mobile elements. Together with previous studies on *P. camemberti, P. roqueforti* and *Geotrichum candidum,* our findings reinforce the view that fungal domestication provides a powerful model for understanding rapid adaptation to human-created environments.

## Material and Methods

### Sampling and DNA extraction

Several strains and genomes were already available (O’Donnell et al., 2025), as indicated in Table S1 (column R). For new strains and genomes, we isolated fungal strains from artisanal and industrial cheeses, contaminated food and other environments. Prior to isolation, all samples were stored at −20 °C for at least 24 hours to inhibit the development of potential mites. Serial dilutions were made and inoculated on Malt Extract Agar (MEA; Sigma-Aldrich, St Louis, MO, USA) or on Rose Bengal Chloramphenicol Agar (RBCA; Difco, Sparks, MD, USA). After five days of incubation at 25°C, distinct morphotypes were isolated. Genomic DNA was extracted using the Nucleospin Soil Kit (Macherey-Nagel, Duren, Germany) and was checked by NanoDrop™ 2000 (Thermofisher Scientific, Waltham, US) and by Qubit™ 4 fluorometer (Thermofisher Scientific, Waltham, US) combined with the Qubit dsDNA BR Assay Kit to estimate if the requirements were reached for the whole genome sequencing. A fragment of β-tubulin gene was amplified by PCR for a 30 μL reactions, using 1 μL of template DNA (100 ng/μL), 0.15 μL of Taq DNA polymerase (5 U/μL, MP Biomedicals), 3 μL of buffer with MgCl_2_ (15mM), 1.2 μL of dNTPs (2.5 mmol/L), 1.2 μL of each primer (bt2a-GGTAACCAAATCGGTGCTGCTTTC and bt2b – AACCTCAGTGTAGTGACCCTTGGC) (Glass and Donaldson, 1995) and 22.5 μL of water. The thermal cycling conditions were as follows: initial denaturation at 94 °C for 5 min; 30 cycles of 94 °C for 30 s, 55 °C for 30 s, and 72 °C for 1 min; followed by a final extension at 72 °C for 5 min. Amplification of a 500bp fragment of β-tubulin gene with bt2a and bt2b primers was verified by electrophoresis on a 1.5% agarose gel. The PCR products were submitted to an external sequencing provider (Azenta Life Science) for Sanger sequencing, and the resulting sequences were compared against the NCBI database using BLASTn to determine species identity.

### Genome sequencing

Short-read sequencing was performed with Illumina NovaSeq X Plus Series (PE150) paired-end technology (Illumina Inc.) with 1G of raw data per sample by the Novogene GmbH company (Munich, Germany)..

Long-read sequencing was performed with the Oxford Nanopore Technology (ONT). We prepared a duplex library using the Native Barcoding Kit 24 V14 (SQK-NBD114.24) following the manufacturer’s instructions starting from 1,5 µg genomic DNA. The library was loaded on a R10.4.1 Flo-Min114 flowcell on a MK1C device (MinKNOW 23.03.12) for a 72 hour run with the Fastcalling basecalling algorithm.

### Short-read assemblies

All Illumina reads were trimmed and adapters cleaned with Trimmomatic v0.39 (Bolger et al., 2014). Leading or trailing low-quality or base pairs below a quality score of three were removed. For each read, only parts that had an average quality score higher than 20 on a four base pair (bp) window are kept. After these steps, only reads with a length of at least 36 bp were kept. Cleaned Illumina reads were assembled with SPAdes v3.15.5 (Prjibelski et al., 2020) not using unpaired reads with “--careful” parameter.

### Long-read assemblies

Data were base called using Guppy (v7.1.4; model dna_r10.4.1_e8.2_400bps_fast@v4.2.0) and assembled using Canu (v2.2) (Koren et al., 2017) with all pass reads and Flye (2.9.3-b1797) (Kolmogorov et al., 2019) after using Filtlong (v0.2.1) (github.com/rrwick/Filtlong) to remove all reads shorter than 5 kb. These assemblies were merged using Ragtag’s (v2.1.0) (Alonge et al., 2022) patch function with the Flye assembly as the “target” and the Canu assembly as the “query”. A second round of merging was done replacing the Canu assembly with the merged assembly from the first round. This merged assembly was then polished with Medaka (v2.0.1) (github.com/nanoporetech/medaka) using all the pass ONT reads. The assembly was then polished using the Illumina reads with Hapo-G (v1.3.8) (Aury and Istace, 2021). Finally contig overlaps had been removed with Purge_dups (v1.2.6) (Guan et al., 2020).

### Annotation

All genomes were annotated. Firstly we used Braker (v3.0.8) (--fungus --gff3) (Gabriel et al., 2024) for genome structure annotation using a set of all BUSCOs in fungal taxonomic groups ((Manni et al., 2021). Four more rounds were then rerun with Braker using the previous run’s augustus protein hints. To assess the quality of the structural annotation, the resulting protein dataset was then analyzed using the fungi_odb10 BUSCO dataset. 99.3% to 99.6% of all proteins were annotated for ONT based assemblies and 99.2% to 99.6% for Illumina based assemblies. The resulting proteins were then functionally annotated using Funannotate (v1.8.15) (Palmer & Stajich, 2020). First, we have generated a funannotate database with the “setup” command (-b fungi). Next, the funannotate iprscan wrapper (Interproscan interproscan-5.56-89.0) (Jones et al., 2014) and eggnog-mapper (v2.1.12) (Cantalapiedra et al., 2021) were used to annotate protein domains and Eggnog functions,respectively. The functional annotations and structural annotation were finally aggregated with the aggregate’s function “annotate”.

### Mapping

Cleaned reads were mapped on the reference genome ESE01543 using Bowtie2 v2.5.1 (Langmead & Salzberg, 2012). Maximum fragment length was set to 1000 and the preset “very-sensitive-local” was used. SAMtools v1.18 (Li et al., 2009) was used to filter out duplicate reads and reads with a mapping quality score above ten for SNP calling.

Single nucleotide polymorphisms (SNPs) were called using GATK v4.4.0 (Auwera & O’Connor, 2020; McKenna et al., 2010) HaplotypeCaller, which provides one gVCF per strain (option - ERC GVCF). GVCFs were combined using GATK CombineGVCFs, genotypes with GATK GenotypeGVCFs, SNPs were selected using GATK SelectVariants (option -select-type SNP). SNPs were filtered using GATK VariantFiltration and options QUAL < 30, DP < 10, QD < 2.0, FS > 60.0, MQ < 40.0, SOR > 3.0, QRankSum < −12.5, ReadPosRankSum < −8.0. All processes from cleaning to variant calling were performed with Snakemake v7.32.4 (Mölder et al., 2021).

### Statistics of population genetics

Nucleotide diversity indices (π and Watterson’s θ) and genetic differentiation indices (*FST*) were calculated using the R package PopGenome (Pfeifer et al., 2014) for *P. fuscoglaucum, P. biforme cheesy* and *natural* and *P. camemberti.* π and Watterson’s θ were calculated per site.

### Genetic structure

#### Neighbor-net analysis

We used the SplitsTree App (v6.4.7) (Huson & Bryant, 2024) to generate the neighbor-net based on SNPs with default options. The input contained 97 taxa of *Penicillium biforme* and *P. camemberti* and 417,650 SNPs.

#### Principal component analysis

We used the R package ade4 (v1.7-22, (Dray & Dufour, 2007)) to perform a principal component analysis based on 97 taxa and 417,650 SNPs.

#### Admixture analysis

We used NGSadmix (Skotte et al., 2013) to infer admixture proportions from SNP data, inferring the number of ancestral populations (K) from 2 to 8.

### *Starship* identification

#### *Starship* and *Starship*-like region characterisation and genotyping

All analysis scripts and outputs can be found at https://github.com/SAMtoBAM/biforme_starships/. *Starships* and *Starship*-like regions (SLRs) were identified using Starfish (Gluck-Thaler & Vogan, 2024) and Stargraph (O’Donnell et al., 2025, 2026). All genomes were analysed initially using the Stargraph *starfish_wrapper* script using default options. Following this, putative *Starships* were manually validated by visually inspecting insertion sites for large contiguous flanking regions. Following this long-read assemblies and their corresponding *Starships*/SLRs were subsetted and used as input to *stargraph* using default options. *Starships* and SLRs from both the short-read and long-read assemblies were recombined and analysed by the Stargraph *cargobay* script with a much stricter threshold for similarity based on k-mer profiles compared to the default value of 0.5 (-c 0.9). This same long and short-read assembly combined dataset was also used to cluster all elements using the same sourmash (Irber et al., 2024) derived Jaccard k-mer similarity with mcl (Van Dongen, 2008) method used in Stargraph. All genomes were screened for these elements using the Starfish *coverage* function (-m paired --aligner minimap2) based on paired-end short read data. The average percentage of coverage was calculated for each strain for each *Starship*/SLR cluster (Table SX_tableUsedForFigure3).

#### *Rattus* cargo-gene functional annotation

The function of *Rattus* genes was initially inferred based on functional annotations from Interproscan (Jones et al., 2014), eggnog-mapper (Cantalapiedra et al., 2021) and Funannotate databases (see Annotation section above) (Palmer & Stajich, 2020). However, as is the case for the majority of *Starships*, several *Rattus* cargo genes had no functional annotation. Therefore, in order to infer the function of those proteins without Interpro or Eggnog annotations, we used a combination of Alphafold3 (Abramson et al., 2024) and Foldseek (van Kempen et al., 2024) to find structurally similar proteins with putative functions. We also used this same process to confirm and better characterise proteins with predicted protein domains. Proteins were first folded using the Alphafold3 server, and the best model (.cif file) used as a search query using the Foldseek server with default options. We then manually looked at proteins with high Foldseek probabilities and TM scores.

The protein g4590 from *Rattus* was folded using AlphaFold3 with a model pTM of 0.85. Next, g4590 was identified by FoldsSeek as similar to numerous proteins in the AFDB50 database from fungal species annotated as ‘Hly-III-related protein’ (e.g. *Penicillium chermesinum* protein A0A9W9TBN1; Foldseek probability: 1.00; e-value: 2.46e-14; TM-Score: 0.91855). Furthermore, in the AFDB-proteome database, Foldseek identified similarity to a Hemolysin-III family protein from Histoplasma capsulatum G186AR (C0NFW8) with a TM-score of 0.79 (Foldseek probability: 1.00; e-value: 2.30e-5) (Figure 4C; FigSX.protein_comps_full_all). We therefore considered the protein g4590 to be a Hemolysin-III-like protein.

The protein g4588 from *Rattus* was folded using AlphaFold3 with a model pTM of 0.63. Next, g4588 was identified by Foldseek as similar to a protein from *Frankineae bacterium* MT45 (A0A1G8NXC9) from the AFDB50 database with a TM-Score of 0.7955 (Foldseek probability: 1; e-value: 2.15e-1) (Figure 4D; FigSX.protein_comps_full_all). This alignment was predicted to cover the A0A1G8NXC9 amino acids 110-208; by inspecting the UniProt database, this portion overlapped with a predicted protein domain superfamily from the SUPFAM database (SSF111369), annotated as a Hemolysin secretion protein D domain. The second-best match from the AFDB50 database was a protein from *Aquibacillus salsiterrae* (A0A9X3WEI7) with a TM-Score of 0.61579 (Foldseek probability: 1.00; e-value: 1.10e-1), which is also assigned to the same Hemolysin-D SUPFAM superfamily. Furthermore, Foldseek identified numerous other proteins from the AFDB50 database, from different bacterial species, that were specifically annotated as ‘HlyD family secretion protein’. We therefore considered the protein g4588 to be a Hemolysin-D-like protein.

The protein g4583 from *Rattus* was folded using AlphaFold3 with a model pTM of 0.71. Next, g4583 was identified by FoldsSeek as similar to numerous proteins in the AFDB50 database from fungal species annotated as ‘Ras-GEF domain-containing protein’ (e.g. *Penicillium daleae* protein A0AAD6C3U5; Foldseek probability: 1.00; e-value: 2.53e-20; TM-Score: 0.6681). Furthermore, in the AFDB-proteome database, Foldseek identified similarity to a Ras-GEF domain-containing protein from *Histoplasma capsulatum* G186AR (C0NLP5) with a TM-score of 0.73134 (Foldseek probability: 1.00; e-value: 4.87e-8) (Figure 4E; FigSX.protein_comps_full_all). We therefore considered the protein g4583 to be a Ras-GEF-like domain protein.

For the two remaining un-annotated *Rattus* proteins, g4587 and g4584, AlphaFold3 did not allow predicting any function, with pTMs of 0.36 and 0.34 respectively, and we did not proceed further.

The protein g4589, annotated as a chitinase (IPR001579, PF00704, IPR011583, IPR017853, IPR001223, IPR029070), was folded using AlphaFold3 with a model pTM of 0.9235. Next, g4589 was identified by Foldseek as similar to numerous chitinases, specifically endochitinases, with the most similar being the ndochitinase B1 from *Aspergillus fumigatus* (E9QRF2; Foldseek probability: 1.00; e-value: 1.20e-71; TM-score: 0.9235) from the AFDB-Swissprot database, considered to be a ‘major secreted chitinase’. Several other matches were also within the Endochitinase B1 CATH Superfamily (3.10.50.10). For example, the ‘endochitinase 1’ protein E9ERT9 from *Metarhizium robertsii* ARSEF 23 (Foldseek probability: 1.00; e-value: 1.28e-66; TM-score 0.93059) (FigSX.protein_comps_full_all). We therefore considered this protein to be an endochitinase.

The protein g4585, annotated as an Allergen Asp f 4 protein (IPR38903), was folded using AlphaFold3 with a model pTM of 0.82. Next, g4585 was identified by Foldseek as similar to numerous Allergen Asp f4 proteins, such as the originally described protein from *Aspergillus fumigatus* (O60024; Foldseek probability: 1.00; e-value: 1.32e-32; TM-score: 0.82559) from the AFDB-Swissprot database. Confirming the structural similarity, the highest e-value for a protein the AFDB-proteome database was an Allergen Asp f4 proteins from *Paracoccidioides lutzii* Pb01 (C1GQR5; Foldseek probability: 1.00; e-value: 7.40e-33; TM-score: 0.89877) (FigSX.protein_comps_full_all). We therefore are confident regarding the annotation for this protein.

Both g4592/Fungal TF and g4586/MADS box TF folded poorly in Alphafold3, with pTM scores of 0.58 and 0.57 respectively, and we did not proceed further.

#### Core and *Rattus* endochitinase gene genealogy

We used the core homologs of the *Rattus* endochitinase in order to determine the cargo genes origin. To do this, we started with the *Rattus* endochitinase from LCP05531. We then expanded this dataset by identifying, using BLASTp (Camacho et al., 2009), all proteins from the annotated genomes that matched the initial subset with at least 60% sequence identity and 50% query coverage. Following this, the protein dataset for each protein was expanded, using the same BLASTp parameters, using a set of publicly available NCBI protein datasets using the NCBI Datasets command-line tools (*datasets download genome accession GCA_XXXXXXXXX* --include protein) (O’Leary et al., 2024). The publicly available protein datasets originated from 24 *Penicillium* and 7 *Aspergillus* genomes, encompassing 15 and 12 species per genus, respectively; this includes the *Pencillium salamii* genome from strain PN007 (GCA911456365) in which *Rattus* was also identified. The final dataset for each *Rattus* protein was therefore a combination of those with the tag in the gff3, and all proteins with BLASTp-based similarity from the annotated proteins of our long-read assemblies and publicly available genomes, including the closest genera as an outgroup. For each protein, amino-acid sequences were aligned using MAFFT (*--auto*) (Katoh & Standley, 2013), alignments trimmed using trimAl(*-automated1*) (Capella-Gutiérrez et al., 2009) and phylogenies built using IQTREE2 (*-B 1000 -alrt 1000*) (Hoang et al., 2018; Kalyaanamoorthy et al., 2017; Minh et al., 2020) and removed branches with less than 50% ultra-fast bootstrapping and/or ALRT support. This tree contained two distinct branches, each with a core endochitinase protein and an *Aspergillus* root. We isolated the clade that contained the *Rattus*-endochitinase and its closest core homolog. These proteins were then used to build a tree, as above, specific to this protein clade (FigSX.endochitinase.phylogeny.full.mod). The set of genomes from which we derived the protein annotations were also used to generate a species tree using Mashtree with bootstrapping (Katz et al., 2019) (*mashtree_bootstrap.pl --reps 1000 *.fna mindepth 0 --sort-order random*). We used both the conservation of endochitinase across multiple genomes of the same species and the genome-derived species tree to identify the core endochitinase homologs.

Using the same BLASTp thresholds as above, the Allergen Asp F 4 was the only other protein in *Rattus* with detectably similarities to proteins in our genome database. We applied the same BLASTp, MAAFT, trimAl, IQTREE2 and Aspergillus rooting as with the endochitinase above. However, the history of non-*Starship* related homologs is more complicated than for the endochitinase genes, with likely duplications (FigSX.AspF4_tree.unrooted.mod). Based on this analysis the originally identified and named protein Allergen Asp F 4 protein from *Aspergillus fumigatus* was detected confirming our ability to detect similar proteins. We manually verified that both proteins in *A. fumigatus* were not in any annotated *Starships* or SLRs using data from a comprehensive Stargraph run (O’Donnell et al., 2026).

### Strain phenotyping

#### Sampling, strain calibration and media used

We chose at random strains of *P. biforme* (n = 20) for phenotypic characterization, with 10 strains of *P. biforme* isolated from cheese and inoculated during cheese production, and 10 strains of *P. biforme* isolated from other environments than food or not intentionally inoculated in cheese. We also chose at random strains of *P. fuscoglaucum* (n = 10) for comparison. The strains were inoculated on MEA; after 5 days of growth, spores were harvested and suspended in sterile water and calibrated to 1.10⁶ spores/mL using a Malassez counting chamber for inoculation of 15 µL on 90mm-diameter petri dishes in all growth experiments detailed below. Each Petri dish was filled with 25mL medium.

Various media were used in this study. Minimal medium (MM), 2% (w/v) agar was prepared according to Hill and Kafer (Hill & Kafer, 2001). Four solutions were prepared. Trace element stock solution was obtained by dissolving FeSO₄·7H₂O (1 g) and EDTA (10 g) in 80 mL of distilled water, followed by pH adjustment above 5.5 using KOH. A second solution containing ZnSO₄·7H₂O (4.4 g), H₃BO₃ (2.2 g), MnCl₂·4H₂O (1 g), CoCl₂·6H₂O (0.32 g), CuSO₄·5H₂O (0.32 g), and (NH₄)₆Mo₇O₂₄·4H₂O (0.22 g) was prepared in 80 mL distilled water. Both solutions were combined, adjusted to pH 6.5, brought to a final volume of 200 mL, and stored at 4°C. A 20× salt stock solution was prepared by dissolving NaNO₃ (120 g), KCl (10.4 g), KH₂PO₄ (16.3 g), and K₂HPO₄ (20.9 g) in distilled water to a final volume of 1 L. A 200× MgSO₄ stock solution was prepared by dissolving MgSO₄·7H₂O (10.4 g) in distilled water to a final volume of 100 mL. For preparation of 1 L of MM, 50 mL of 20× salt stock, 5 mL of 200× MgSO₄ stock, 10 g glucose, and 1 mL trace element solution were added to 950 mL distilled water with agar at 2%.

Cheese medium 1.5% (w/v) NaCl %, 2% (w/v) agar was prepared using raw goat curdled milk obtained from La Doudou Farm (Cheptainville, 91630, France). Defrosted cheese curdled milk (300g) was homogenized in 200 mL of distilled water, the pH was measured and adjusted to 6. A NaCl solution (1.5%, w/v) was prepared in 200 mL of distilled water and combined with the cheese mixture. The final pH was adjusted to 6.0, and the final volume was brought to 1 L with distilled water with 2% agar.

Malt Extract Agar (MEA), 2% (w/v) agar was prepared by dissolving 20 g of malt extract (Sigma-Aldrich) and 20 g of agar in distilled water to a final volume of 1 L. YES 1.5% (w/v) agar medium was prepared according to Gillot et al., 2017 and adjusted to pH 4.0. A concentrated 2X solution was prepared in a final volume of 500 mL distilled water containing: sucrose (150 g), yeast extract (10 g), MgSO₄·7H₂O (0.5 g), ZnSO₄·H₂O (0.006 g), CuSO₄·5H₂O (0.005 g) and agar (15 g). Then two solutions were prepared: 0.2M Na₂HPO₄ (28.32 g) and 0.1M citric acid (21 g). All three solutions were sterilized. A phosphate-citrate buffer (pH 4) was prepared by combining 193 mL of 0.2M Na₂HPO₄ solution and 307 mL of 0.1M citric acid solution. This buffer was then appropriately mixed with the 2X YES concentrated solution and poured into Petri dishes.

MM and MEA media were sterilized at 121°C for 15 minutes, YES media and buffers were sterilized at 121°C for 10 minutes, and cheese media at 110°C for 15 minutes in order to avoid caramelization.

Images of the Petri dishes were obtained with a Scan 1200 from Interscience and analysed with Image J software (v1.54g, NIH, USA), by calculating the area of colonies in pixel². Diameters were measured by hand only for the competition assay because they were not visible using images.

#### Measurement of radial growth on MM, cheese-based and MEA at 12°C and 25°C

To compare growth rates between populations, we used three different media: cheese with 1.5% (w/v) NaCl, MEA, rich in carbohydrates and MM which contained basic elements for fungal survival. We incubated Petri dishes at two different conditions: 12°C (cave condition) and 25°C. Images were taken at 7, 14 and 21 days.

#### Measurement of radial growth on cheese-based media supplemented with varying salt concentrations (salt tolerance)

We prepared modified cheese media with five different salt concentrations: 0, 1.5%, 2%, 4% and 8% (w/v). Petri dishes were incubated at 12°C and images were taken weekly across 21 days.

#### Measurement of radial growth on different carbon source media

We prepared modified MM by replacing glucose (55.5mM) as the original carbon source, with lactose or acid lactic or galactose or xylose, keeping 55.5mM as molecular weight for each. Petri dishes were incubated at 25°C and images were taken weekly across 21 days.

#### Measurement of vertical growth on a cheese-based medium (fluffiness)

Six *P. camemberti* strains were included for this experiment. Vertical growth was measured on cheese medium using glass tubes filled with ten milliliters of medium. We inoculated 10 µL of the calibrated spore solutions and tubes were incubated at 12°C for 4 weeks to mimic cave conditions. The fluffiness of each strain, i.e. the vertical growth of the mycelium, was measured by hand with a ruler every 7 days for 28 days.

#### Measurement of lipolytic and proteolytic activities

We inoculated 10 µL of calibrated spore solutions (1.10⁶ spores.mL^-1^) of *P. biforme* (n= 20) and *P. fuscoglaucum* (n=10) in sterilized glass tubes containing agar and semi skimmed milk for proteolysis (40 g/L, skimmed milk Regilait) and agar tributyrin (20 mL.L^-1^, ACROS Organics) for lipolysis (Dumas, et al, 2020). The semi-skimmed milk agar was sterilized at 110°C for 15 minutes and the agar tributyrin was sterilized at 121°C for 15 minutes. The tubes were incubated at 12°C and the degradation of compounds was measured by hand with a ruler according to the change in opacity to a translucent medium after 7, 14, 21 and 28 days.

#### Measurement of spore production and colony count

After 24 days of strain growth on MEA, cheese media and MM at 12°C, a mycelial plug was taken from each Petri dish using a 15 mL Falcon tube and transferred to an Eppendorf tube containing sterile water. To determine the concentration of spores, 5 µL of each suspension was diluted in 1:200 and counted using a Malassez counting chamber. To assess colony formation capacity following growth on the different media, the spore concentrations were adjusted to 1000 spores/mL and 100 µL was spread onto MEA Petri dishes. We incubated Petri dishes at 12°C for 5 days, after which we counted the colony forming units (CFU).

#### Volatile organic compound analysis

To assess volatile organic compound (VOC) production, we used a cheese medium supplemented with 1.5% NaCl (w/v). For this analysis, we additionally included strains from *P. camemberti*, as VOC production had not previously been assessed in this group. The sampling comprised *P. camemberti* (n = 3), *P. caseifulvum* (n = 3), *P. biforme* (n = 20), and *P. fuscoglaucum* (n = 4). Thirty milliliters of medium were poured into glass Petri dishes, and each dish was inoculated with 15 µL of a calibrated spore suspension containing 1.10⁶ spores.mL^-1^. Petri dishes were incubated at 12°C for 4 weeks to mimic cheese-ripening conditions.

VOCs were collected for 24h using polydimethylsiloxane (PDMS)-coated stir bars (“Twisters”; Gerstel GmbH, Mülheim an der Ruhr, Germany), following Guo et al. (Guo et al., 2021). Subsequently, VOCs were analyzed using a GERSTEL Thermal Desorption Unit (TDU), which was coupled with a Cooled Injection System (CIS 4) and an Agilent (Agilent, Waldbronn, Germany) 7890A gas chromatograph (GC), connected to an Agilent 5975C mass selective detector (MSD). Each Twister spiked with 1 µL of internal Standard (ISTD: δ-2-carene solved in hexane; c=859.28 pmol.µL^-1^) before measurement. After ISTD applied each twister was thermally desorbed in the TDU using the following program: an initial temperature of 30 °C (initial time 0.10 min), ramped at a rate of 280 °C.min^-1^ to 270 °C (held for 2 min). The desorbed compounds were cryo focused on a glass liner filled with tenax and glass wool in the CIS at −50 °C, the transferline between TDU and CIS was at a constant 250°C. Following an additional desorption step after 0.31min in the CIS with 12°C.sec^-1^ to an end temperature of 270°C which was held for 2 min under a constant flow of 104 mL.min^-1^ which transfers the sample directly onto the GC-Column in splitless mode. Gas chromatographic separation was performed on a 60m DB-5MS + 10m DG (5%-phenyl)-methylpolysiloxane column (70 m × 0.25 mm i.d. × 0.25 µm film thickness; Agilent J&W 122-5562G) using helium 5.0 as the carrier gas at a constant flow rate of 1.0 mL min-1. The GC oven temperature program started with an initial temperature of 40 °C, followed by ramping at 10 °C min⁻¹ to 130 °C (held for 5 min), then at 80 °C min⁻¹ to 175 °C, then at 2 °C min⁻¹ to 200 °C, then at 4 °C min⁻¹ to 220 °C, and finally at 100 °C min⁻¹ to 300 °C (held for 5 min). The total run time was 38.86 minutes. The MSD operated in electron-impact (EI) mode at 70 eV. The MS source and quadrupole temperatures were maintained at 230 °C and 150 °C, respectively. Data was acquired in combined SIM/scan mode after a solvent delay of 8.0 minutes. Full scan mode monitored a mass range of m/z 35-250. To enhance sensitivity for key compound classes, two SIM groups were employed: one for monoterpenes (m/z ions 93, 121 and 136), which started at 8.0 minutes, and one for sesquiterpenes (m/z ions 93, 161 and 204), which started at 14.9 minutes. Compounds were identified by comparing their mass spectra in the NIST 2020 spectral library (match quality >70%). Quantification was performed using external calibration curves. Potential changes in GC-MS sensitivity were corrected by normalization to the internal standard (Ghirardo et al., 2011, 2020).

For biological interpretation, VOCs were curated and assigned to aroma-relevant sensory families. Styrene was excluded, as is standard practice, due to its common association with laboratory plastic materials, packaging components or laboratory-derived contamination, which precludes confident attribution to fungal metabolism. Additionally, four unidentified VOC features, annotated as cyclic-aldehyde_01–04, were excluded because they lacked validated odor descriptors. While these compounds were retained in the raw analytical dataset, they were omitted from the aroma-family assignment.

The remaining VOCs were classified into seven aroma families based on odor descriptors sourced from two public databases ((https://www.thegoodscentscompany.com/ and https://scent.vn/en). The defined aroma families were dairy/cheesy, mushroom, fatty, fruity, woody/spicy/herbal, plastic, aldehydic/unknown.

#### Mycotoxin production

We only assessed mycotoxin production for *P. biforme* strains (n = 30) as *P. camemberti* lineages and *P. fuscoglaucum* strains were previously screened (Ropars et al., 2020). Each strain was grown on YES agar using six Petri dishes, three with sterilized cellulose (to facilitate mycelium removal and determine dry weights for normalizing toxin production) and three without sterilized cellulose (to quantify toxin production) to ensure triplicates for each strain. We inoculated 1 µL (1.10⁶ spores.mL^-1^) of each *P. biforme* strain in the center, and incubated Petri dishes at 25°C for 10 days. Mycelium was dried after growth on cellulose for 4 days at 70°C and fungal dry weight was measured. For mycotoxin determination, fungal strains on YES agar without cellulose were collected in 50 mL Falcon tubes and stored at −80°C until mycotoxin extraction and quantification.

For mycotoxin extractions, we used an optimised high-throughput extraction method (Crequer et al., 2024; Gillot et al., 2017; Lo et al., 2023). Briefly, samples were thawed before being homogenised until a homogenous mixture was obtained. We then sampled 2 g aliquots and added 12.5 mL of acetonitrile (ACN) supplemented with 0.1% formic acid (v/v); samples were vortexed followed by 15 min sonication. The extracts were vortexed again before centrifugation for 10 min at 5,000 *g* at 4 °C. The supernatants were then collected and filtered through 0.45 μm polytetrafluoroethylene membrane filters into amber vials and stored at −20 °C until analysis.

Cylcopiazonic acid (CPA) standard was obtained from Sigma-Aldrich (St Louis, MO, USA). A stock solution was prepared in dimethyl sulfoxide (DMSO) at 1 mg.mL^-1^ in amber vials. For CPA quantification, toxin identification was performed using both the mean retention time ± 1 min and the corresponding ion ([M+H]+ 337.1546) in electrospray ionization mode positive (ESI+). We used a matrix-matched calibration curve to ensure reliable mycotoxin quantification with final concentrations ranging from 500 to 10000 ng.mL−1; method performance was as previously described (Gillot et al. 2017) and all quantification analyses were determined using the Agilent MassHunter Workstation Software (Agilent Technologies, Sanat Clara, CA, USA) with a linear regression model. Specific mycotoxin production was expressed as ng per g of fungal dry weight (ng.g^-1^). Linearity (R2) was determined to be 0.997 in ESI +. CPA analyses were performed on an Agilent 6530 Accurate-Mass Quadrupole Time-of-Flight mass spectrometry system equipped with a binary pump 1260 and degasser (Q-TOF LC/MS), well plate autosampler set to 10 °C and a thermo-stated column compartment. Filtered 2 μL aliquots were injected into a ZORBAX Extend C-18 column (2.1 × 50 mm and 1.8 μm, 600 bar) maintained at 35 °C with a flow rate set to 0.3 mL.min^-1^. The mobile phase A contained milli-Q water + 0.1% formic acid (v/v) while mobile phase B was ACN + 0.1% formic acid. Mobile phase B was maintained at 10% for 4 min followed by a gradient from 10 to 100% for 15 min. Then, mobile phase B was maintained at 100% for 5 min before a 5 min post-time. Samples were ionised in ESI+ mode in the mass spectrometer with the following parameters: capillary voltage 4 kV, source temperature 325 °C, nebulizer pressure 50 psig, drying gas 12 L.min^-1^, ion range 100–1000 m/z.

#### Competition assays

We assessed contaminant exclusion capacity on cheese medium with 1.5% (w/v) NaCl by growing mycelium lawns of tested strains and inoculating them with challengers. We also assessed the inhibition capacity through volatile compounds by growing tested strains and challengers in separated compartments of a single Petri dish (plastic physical barrier). We selected four species of challengers to test each against our 30 *P. biforme* and *P. fuscoglaucum* strains: *Yarrowia lipolytica* (n=5), *Scopulariopsis asperula* (n=5)*, Mucor racemosus* (n=5)*, Penicillium fuscoglaucum* (n=5).

| Strain (ESE collection) | Species | Isolation |
| --- | --- | --- |
| ESE00273 | *Yarrowia lipolytica* | Neufchâtel (GAEC Fouque) |
| ESE00277 | *Yarrowia lipolytica* | Pont l'èvêque Graindorge (industrial) |
| ESE00281 | *Yarrowia lipolytica* | Ganot farm (Seine et Marne) |
| ESE00964 | *Yarrowia lipolytica* | Gamonio Astorias (Spain) |
| ESE01386 | *Yarrowia lipolytica* | Ossau-Iraty cheese (artisanal, Pyrénées) |
| ESE01324 | *Scopulariopsis asperula* | Ossau Iraty (Pyrenees) |
| ESE01463 | *Scopulariopsis asperula* | Tomme goat cheese |
| ESE01577 | *Scopulariopsis asperula* | Tomme cow Savoie |
| ESE01582 | *Scopulariopsis asperula* | Tomme ewe |
| ESE01614 | *Scopulariopsis asperula* | Tome du Berry |
| ESE01882 | *Mucor racemosus* | Cheese |
| ESE01883 | *Mucor racemosus* | Cheese rind |
| ESE01884 | *Mucor racemosus* | Hard cheese |
| ESE01885 | *Mucor racemosus* | Yogurt |
| ESE01886 | *Mucor racemosus* | Cheese surface |
| ESE00159 | *Penicillium fuscoglaucum* | Cantal |
| ESE00209 | *Penicillium fuscoglaucum* | Insect |
| ESE00216 | *Penicillium fuscoglaucum* | Walnut |
| ESE00237 | *Penicillium fuscoglaucum* | Saint nectaire |
| ESE00266 | *Penicillium fuscoglaucum* | Cheese |

We poured a total of 1240 Petri dishes (90 mm **×** 15 mm): 620 regular Petri dishes and 620 Petri dishes with two compartments separated by a division along a diameter. For assessing contaminant exclusion capacity by mycelium lawns, 100µL of calibrated spore solutions of *P. biforme* or *P. fuscoglaucum* were inoculated and spread on the Petri dishes. After 24h of growth, 5µL of a competitor spore solution (1.10^6^ spores.mL^-1^) were inoculated in a single spot in the middle of each Petri dish. To compare the abilities of the two *P. biforme* populations and *P. fuscoglaucum* to exclude other microorganisms by producing volatile compounds, 50µL of spore solution were spread on one compartment of each Petri dish separated in halves. After 24h of growth, 2.5µL, a competitor spore solution (1.10^6^ spores.mL^-1^) were inoculated in a single spot on the other half of each Petri dish. All the 1,240 plates were incubated at 12°C. The contaminant colony diameters (1 measure per Petri dish) were measured with a ruler at different days following the inoculation: 4 days for *Mucor racemosus,* 7 days for the white *Penicillium fuscoglaucum,* 10 days for S*copulariopsis asperula* and 18 days for *Yarrowia lipolytica*.

## Data and scripts availability

Data and scripts are available online. New genome sequencing and assemblies have been deposited on NCBI (bioproject XX).

## Author contributions

S.O: Data Curation, Formal Analysis, Investigation, Methodology, Software, Validation, Visualization, Writing and editing. GR.: Investigation, Formal analyses on phenotypic tests, visualization, Methodology, writing. J.-P.V.: Data Curation, Investigation, Software. A.S.: Investigation. A.L.: Investigation. M.C.: secondary metabolites analyses. J.-P.S.: Analyses of volatiles, writing-review. B.W.: Investigation and analyses of volatiles. T.G.: Writing-review. J.R.: Conceptualization, Data Curation, Funding acquisition, Formal Anaysis, Investigation, Methodology, Project administration, Resources, Supervision, Validation, Writing-original draft, Writing-review and editing.

## Supporting information

Strain information

## References

Abramson, J., Adler, J., Dunger, J., Evans, R., Green, T., Pritzel, A., Ronneberger, O., Willmore, L., Ballard, A. J., Bambrick, J., Bodenstein, S. W., Evans, D. A., Hung, C.-C., O’Neill, M., Reiman, D., Tunyasuvunakool, K., Wu, Z., Žemgulytė, A., Arvaniti, E., … Jumper, J. M. (2024). Accurate structure prediction of biomolecular interactions with AlphaFold 3. Nature, 630(8016), 493–500. 10.1038/s41586-024-07487-w

Alonge, M., Lebeigle, L., Kirsche, M., Jenike, K., Ou, S., Aganezov, S., Wang, X., Lippman, Z. B., Schatz, M. C., & Soyk, S. (2022). Automated assembly scaffolding using RagTag elevates a new tomato system for high-throughput genome editing. Genome Biology, 23(1), 258. 10.1186/s13059-022-02823-7

Arnold, B. J., Huang, I.-T., & Hanage, W. P. (2022). Horizontal gene transfer and adaptive evolution in bacteria. Nature Reviews Microbiology, 20(4), 206–218. 10.1038/s41579-021-00650-4

Auwera, G. A. V. der, & O’Connor, B. D. (2020). Genomics in the Cloud: Using Docker, GATK, and WDL in Terra. O’Reilly Media, Inc.

Bennetot, B., Vernadet, J.-P., Perkins, V., Hautefeuille, S., Rodríguez De La Vega, R. C., O’Donnell, S., Snirc, A., Grondin, C., Lessard, M.-H., Peron, A.-C., Labrie, S., Landaud, S., Giraud, T., & Ropars, J. (2023). Domestication of different varieties in the cheese-making fungus Geotrichum candidum. Peer Community Journal, 3, e45. 10.24072/pcjournal.266

Bertuzzi, A. S., McSweeney, P. L. H., Rea, M. C., & Kilcawley, K. N. (2018). Detection of Volatile Compounds of Cheese and Their Contribution to the Flavor Profile of Surface-Ripened Cheese. Comprehensive Reviews in Food Science and Food Safety, 17(2), 371–390. 10.1111/1541-4337.12332

Bolger, A. M., Lohse, M., & Usadel, B. (2014). Trimmomatic: A flexible trimmer for Illumina sequence data. Bioinformatics (Oxford, England), 30(15), 2114–2120. 10.1093/bioinformatics/btu170

Bucknell, A., Wilson, H. M., Gonçalves dos Santos, K. C., Simpfendorfer, S., Milgate, A., Germain, H., Solomon, P. S., Bentham, A., & McDonald, M. C. (2025). Sanctuary: A Starship transposon facilitating the movement of the virulence factor ToxA in fungal wheat pathogens. mBio, 0(0), e01371–25. 10.1128/mbio.01371-25

Camacho, C., Coulouris, G., Avagyan, V., Ma, N., Papadopoulos, J., Bealer, K., & Madden, T. L. (2009). BLAST+: Architecture and applications. BMC Bioinformatics, 10(1), 421. 10.1186/1471-2105-10-421

Cantalapiedra, C. P., Hernández-Plaza, A., Letunic, I., Bork, P., & Huerta-Cepas, J. (2021). eggNOG-mapper v2: Functional Annotation, Orthology Assignments, and Domain Prediction at the Metagenomic Scale. Molecular Biology and Evolution, 38(12), 5825–5829. 10.1093/molbev/msab293

Capella-Gutiérrez, S., Silla-Martínez, J. M., & Gabaldón, T. (2009). trimAl: A tool for automated alignment trimming in large-scale phylogenetic analyses. Bioinformatics, 25(15), 1972–1973. 10.1093/bioinformatics/btp348

Caron, T., Piver, M. L., Péron, A.-C., Lieben, P., Lavigne, R., Brunel, S., Roueyre, D., Place, M., Bonnarme, P., Giraud, T., Branca, A., Landaud, S., & Chassard, C. (2021). Strong effect of Penicillium roqueforti populations on volatile and metabolic compounds responsible for aromas, flavor and texture in blue cheeses. International Journal of Food Microbiology, 354, 109174. 10.1016/j.ijfoodmicro.2021.109174

Cheeseman, K., Ropars, J., Renault, P., Dupont, J., Gouzy, J., Branca, A., Abraham, A.-L., Ceppi, M., Conseiller, E., Debuchy, R., Malagnac, F., Goarin, A., Silar, P., Lacoste, S., Sallet, E., Bensimon, A., Giraud, T., & Brygoo, Y. (2014). Multiple recent horizontal transfers of a large genomic region in cheese making fungi. Nature Communications, 5, 2876. 10.1038/ncomms3876

Crequer, E., Coton, E., Cueff, G., Cristiansen, J. V., Frisvad, J. C., Rodríguez de la Vega, R. C., Giraud, T., Jany, J.-L., & Coton, M. (2024). Different metabolite profiles across Penicillium roqueforti populations associated with ecological niche specialisation and domestication. IMA Fungus, 15(1), 38. 10.1186/s43008-024-00167-4

Crequer, E., Ropars, J., Jany, J.-L., Caron, T., Coton, M., Snirc, A., Vernadet, J.-P., Branca, A., Giraud, T., & Coton, E. (2023). A new cheese population in *Penicillium roqueforti* and adaptation of the five populations to their ecological niche. Evolutionary Applications, 00, 1–20. 10.1111/eva.13578

Delfosse, C. (2008). Histoires de bries. Illustria Librairie des Musées.

Dray, S., & Dufour, A.-B. (2007). The ade4 Package: Implementing the Duality Diagram for Ecologists. Journal of Statistical Software, 22, 1–20. 10.18637/jss.v022.i04

Dumas, E., Feurtey, A., Vega, R. C. R. de la, Prieur, S. L., Snirc, A., Coton, M., Thierry, A., Coton, E., Piver, M. L., Roueyre, D., Ropars, J., Branca, A., & Giraud, T. (2020). Independent domestication events in the blue-cheese fungus *Penicillium roqueforti*. Molecular Ecology, 29(14), 2639–2660. 10.1111/mec.15359

Etten, J. V., & Bhattacharya, D. (2020). Horizontal Gene Transfer in Eukaryotes: Not if, but How Much? Trends in Genetics, 36(12), 915–925. 10.1016/j.tig.2020.08.006

Gabriel, L., Brůna, T., Hoff, K. J., Ebel, M., Lomsadze, A., Borodovsky, M., & Stanke, M. (2024). BRAKER3: Fully automated genome annotation using RNA-seq and protein evidence with GeneMark-ETP, AUGUSTUS, and TSEBRA. Genome Research, 34(5), 769–777. 10.1101/gr.278090.123

Ghirardo, A., Gutknecht, J., Zimmer, I., Brüggemann, N., & Schnitzler, J.-P. (2011). Biogenic Volatile Organic Compound and Respiratory CO2 Emissions after 13C-Labeling: Online Tracing of C Translocation Dynamics in Poplar Plants. PLOS ONE, 6(2), e17393. 10.1371/journal.pone.0017393

Ghirardo, A., Lindstein, F., Koch, K., Buegger, F., Schloter, M., Albert, A., Michelsen, A., Winkler, J. B., Schnitzler, J.-P., & Rinnan, R. (2020). Origin of volatile organic compound emissions from subarctic tundra under global warming. Global Change Biology, 26(3), 1908–1925. 10.1111/gcb.14935

Gillot, G., Jany, J.-L., Poirier, E., Maillard, M.-B., Debaets, S., Thierry, A., Coton, E., & Coton, M. (2017). Functional diversity within the *Penicillium roqueforti* species. International Journal of Food Microbiology, 241, 141–150. 10.1016/j.ijfoodmicro.2016.10.001

Gladieux, P., Ropars, J., Badouin, H., Branca, A., Aguileta, G., de Vienne, D. M., Rodríguez de la Vega, R. C., Branco, S., & Giraud, T. (2014). Fungal evolutionary genomics provides insight into the mechanisms of adaptive divergence in eukaryotes. Molecular Ecology, 23(4), 753–773. 10.1111/mec.12631

Gluck-Thaler, E., Forsythe, A., Puerner, C., Gutierrez-Perez, C., Stajich, J. E., Croll, D., Cramer, R. A., & Vogan, A. A. (2025). Giant transposons promote strain heterogeneity in a major fungal pathogen. mBio, 16(6), e01092–25. 10.1128/mbio.01092-25

Gluck-Thaler, E., & Vogan, A. A. (2024). Systematic identification of cargo-mobilizing genetic elements reveals new dimensions of eukaryotic diversity. Nucleic Acids Research, 52(10), 5496–5513. 10.1093/nar/gkae327

Guan, D., McCarthy, S. A., Wood, J., Howe, K., Wang, Y., & Durbin, R. (2020). Identifying and removing haplotypic duplication in primary genome assemblies. Bioinformatics, 36(9), 2896–2898. 10.1093/bioinformatics/btaa025

Guo, Y., Jud, W., Weikl, F., Ghirardo, A., Junker, R. R., Polle, A., Benz, J. P., Pritsch, K., Schnitzler, J.-P., & Rosenkranz, M. (2021). Volatile organic compound patterns predict fungal trophic mode and lifestyle. Communications Biology, 4(1), Article 1. 10.1038/s42003-021-02198-8

Hartmann, F. E., Rodríguez de la Vega, R. C., Demené, A., Badet, T., Vernadet, J.-P., Rougemont, Q., Labat, A., Snirc, A., Stauber, L., Croll, D., Prospero, S., Dutech, C., & Giraud, T. (2025). An Inversion Polymorphism Under Balancing Selection, Involving Giant Mobile Elements, in an Invasive Fungal Pathogen. Molecular Biology and Evolution, 42(2), msaf026. 10.1093/molbev/msaf026

Hill, T. W., & Kafer, E. (2001). Improved protocols for Aspergillus minimal medium: Trace element and minimal medium salt stock solutions. Fungal Genetics Reports, 48(1), 20–21. 10.4148/1941-4765.1173

Hoang, D. T., Chernomor, O., von Haeseler, A., Minh, B. Q., & Vinh, L. S. (2018). UFBoot2: Improving the Ultrafast Bootstrap Approximation. Molecular Biology and Evolution, 35(2), 518–522. 10.1093/molbev/msx281

Huson, D. H., & Bryant, D. (2024). The SplitsTree App: Interactive analysis and visualization using phylogenetic trees and networks. Nature Methods, 21(10), 1773–1774. 10.1038/s41592-024-02406-3

Irber, L., Pierce-Ward, N. T., Abuelanin, M., Alexander, H., Anant, A., Barve, K., Baumler, C., Botvinnik, O., Brooks, P., Dsouza, D., Gautier, L., Hera, M. R., Houts, H. E., Johnson, L. K., Klötzl, F., Koslicki, D., Lim, M., Lim, R., Nelson, B., … Brown, C. T. (2024). sourmash v4: A multitool to quickly search, compare, and analyze genomic and metagenomic data sets. Journal of Open Source Software, 9(98), 6830. 10.21105/joss.06830

Irlinger, F., Mariadassou, M., Dugat-Bony, E., Rué, O., Neuvéglise, C., Renault, P., Rifa, E., Theil, S., Loux, V., Cruaud, C., Gavory, F., Barbe, V., Lasbleiz, R., Gaucheron, F., Spelle, C., & Delbès, C. (2024). A comprehensive, large-scale analysis of “terroir” cheese and milk microbiota reveals profiles strongly shaped by both geographical and human factors. ISME Communications, 4(1), ycae095. 10.1093/ismeco/ycae095

Jones, P., Binns, D., Chang, H.-Y., Fraser, M., Li, W., McAnulla, C., McWilliam, H., Maslen, J., Mitchell, A., Nuka, G., Pesseat, S., Quinn, A. F., Sangrador-Vegas, A., Scheremetjew, M., Yong, S.-Y., Lopez, R., & Hunter, S. (2014). InterProScan 5: Genome-scale protein function classification. Bioinformatics, 30(9), 1236–1240. 10.1093/bioinformatics/btu031

Kalyaanamoorthy, S., Minh, B. Q., Wong, T. K. F., von Haeseler, A., & Jermiin, L. S. (2017). ModelFinder: Fast model selection for accurate phylogenetic estimates. Nature Methods, 14(6), Article 6. 10.1038/nmeth.4285

Karahadian, C., Josephson, D. B., & Lindsay, R. C. (1985). Contribution of Penicillium sp. To the Flavors of Brie and Camembert Cheese1. Journal of Dairy Science, 68(8), 1865–1877. 10.3168/jds.S0022-0302(85)81043-2

Katoh, K., & Standley, D. M. (2013). MAFFT Multiple Sequence Alignment Software Version 7: Improvements in Performance and Usability. Molecular Biology and Evolution, 30(4), 772–780. 10.1093/molbev/mst010

Katz, L. S., Griswold, T., Morrison, S. S., Caravas, J. A., Zhang, S., Bakker, H. C. den, Deng, X., & Carleton, H. A. (2019). Mashtree: A rapid comparison of whole genome sequence files. Journal of Open Source Software, 4(44), 1762. 10.21105/joss.01762

Kolmogorov, M., Yuan, J., Lin, Y., & Pevzner, P. A. (2019). Assembly of long, error-prone reads using repeat graphs. Nature Biotechnology, 37(5), 540–546. 10.1038/s41587-019-0072-8

Koren, S., Walenz, B. P., Berlin, K., Miller, J. R., Bergman, N. H., & Phillippy, A. M. (2017). Canu: Scalable and accurate long-read assembly via adaptive k-mer weighting and repeat separation. Genome Research, 27(5), 722–736. 10.1101/gr.215087.116

Langmead, B., & Salzberg, S. L. (2012). Fast gapped-read alignment with Bowtie 2. Nature Methods, 9(4), 357–359. 10.1038/nmeth.1923

Le Bars, J. (1979). Cyclopiazonic acid production by *Penicillium camemberti* Thom and natural occurrence of this mycotoxin in cheese. Applied and Environmental Microbiology, 38(6), 1052–1055.

Li, H., Handsaker, B., Wysoker, A., Fennell, T., Ruan, J., Homer, N., Marth, G., Abecasis, G., Durbin, R., & 1000 Genome Project Data Processing Subgroup. (2009). The Sequence Alignment/Map format and SAMtools. Bioinformatics (Oxford, England), 25(16), 2078–2079. 10.1093/bioinformatics/btp352

Lo, Y., Bruxaux, J., Rodríguez de la Vega, R. C., O’Donnell, S., Snirc, A., Coton, M., Le Piver, M., Le Prieur, S., Roueyre, D., Dupont, J., Houbraken, J., Debuchy, R., Ropars, J., Giraud, T., & Branca, A. (2023). Domestication in dry-cured meat Penicillium fungi: Convergent specific phenotypes and horizontal gene transfers without strong genetic subdivision. Evolutionary Applications, 16(9), 1637–1660. 10.1111/eva.13591

Manni, M., Berkeley, M. R., Seppey, M., Simão, F. A., & Zdobnov, E. M. (2021). BUSCO Update: Novel and Streamlined Workflows along with Broader and Deeper Phylogenetic Coverage for Scoring of Eukaryotic, Prokaryotic, and Viral Genomes. Molecular Biology and Evolution, 38(10), 4647–4654. 10.1093/molbev/msab199

McKenna, A., Hanna, M., Banks, E., Sivachenko, A., Cibulskis, K., Kernytsky, A., Garimella, K., Altshuler, D., Gabriel, S., Daly, M., & DePristo, M. A. (2010). The Genome Analysis Toolkit: A MapReduce framework for analyzing next-generation DNA sequencing data. Genome Research, 20(9), 1297–1303. 10.1101/gr.107524.110

Minh, B. Q., Schmidt, H. A., Chernomor, O., Schrempf, D., Woodhams, M. D., von Haeseler, A., & Lanfear, R. (2020). IQ-TREE 2: New Models and Efficient Methods for Phylogenetic Inference in the Genomic Era. Molecular Biology and Evolution, 37(5), 1530–1534. 10.1093/molbev/msaa015

Mölder, F., Jablonski, K. P., Letcher, B., Hall, M. B., van Dyken, P. C., Tomkins-Tinch, C. H., Sochat, V., Forster, J., Vieira, F. G., Meesters, C., Lee, S., Twardziok, S. O., Kanitz, A., VanCampen, J., Malladi, V., Wilm, A., Holtgrewe, M., Rahmann, S., Nahnsen, S., & Köster, J. (2021). Sustainable data analysis with Snakemake. F1000Research, 10, 33. 10.12688/f1000research.29032.3

Molimard, P., & Spinnler, H. E. (1996). Review: Compounds Involved in the Flavor of Surface Mold-Ripened Cheeses: Origins and Properties. Journal of Dairy Science, 79(2), 169–184. 10.3168/jds.S0022-0302(96)76348-8

O’Donnell, S., McVey, A., Valent, B., Liu, S., Gluck-Thaler, E., & Cook, D. E. (2026). *Distinct modes of sequence evolution and epigenetic modifications underpin the origins of Starship-mediated variation in Pyricularia fungal plant pathogens* (p. 2026.01.28.702382). bioRxiv. 10.64898/2026.01.28.702382

O’Donnell, S., Rezende, G., Vernadet, J.-P., Snirc, A., & Ropars, J. (2025). Harbouring Starships: The accumulation of large Horizontal Gene Transfers in Domesticated and Pathogenic Fungi. *Genome Biology and Evolution*, evaf125. 10.1093/gbe/evaf125

O’Leary, N. A., Cox, E., Holmes, J. B., Anderson, W. R., Falk, R., Hem, V., Tsuchiya, M. T. N., Schuler, G. D., Zhang, X., Torcivia, J., Ketter, A., Breen, L., Cothran, J., Bajwa, H., Tinne, J., Meric, P. A., Hlavina, W., & Schneider, V. A. (2024). Exploring and retrieving sequence and metadata for species across the tree of life with NCBI Datasets. Scientific Data, 11(1), 732. 10.1038/s41597-024-03571-y

Palmer, J. M., & Stajich, J. (2020). *Funannotate v1.8.1: Eukaryotic genome annotation* [Computer software]. Zenodo. 10.5281/zenodo.4054262

Peck, L. D., Llewellyn, T., Bennetot, B., O’Donnell, S., Nowell, R. W., Ryan, M. J., Flood, J., Vega, R. C. R. de la, Ropars, J., Giraud, T., Spanu, P. D., & Barraclough, T. G. (2024). Horizontal transfers between fungal Fusarium species contributed to successive outbreaks of coffee wilt disease. PLOS Biology, 22(12), e3002480. 10.1371/journal.pbio.3002480

Penland, M., Falentin, H., Parayre, S., Pawtowski, A., Maillard, M.-B., Thierry, A., Mounier, J., Coton, M., & Deutsch, S.-M. (2021). Linking Pélardon artisanal goat cheese microbial communities to aroma compounds during cheese-making and ripening. International Journal of Food Microbiology, 345, 109130. 10.1016/j.ijfoodmicro.2021.109130

Pfeifer, B., Wittelsbürger, U., Ramos-Onsins, S. E., & Lercher, M. J. (2014). PopGenome: An Efficient Swiss Army Knife for Population Genomic Analyses in R. Molecular Biology and Evolution, 31(7), 1929–1936. 10.1093/molbev/msu136

Prjibelski, A., Antipov, D., Meleshko, D., Lapidus, A., & Korobeynikov, A. (2020). Using SPAdes De Novo Assembler. Current Protocols in Bioinformatics, 70(1), e102. 10.1002/cpbi.102

Ropars, J., Didiot, E., Rodríguez de la Vega, R. C., Bennetot, B., Coton, M., Poirier, E., Coton, E., Snirc, A., Le Prieur, S., & Giraud, T. (2020). Domestication of the emblematic white cheese-making fungus *Penicillium camemberti* and its diversification into two varieties. Current Biology, 30(22), 4441–4453. 10.1016/j.cub.2020.08.082

Ropars, J., Rodríguez de la Vega, R. C., López-Villavicencio, M., Gouzy, J., Sallet, E., Dumas, É., Lacoste, S., Debuchy, R., Dupont, J., Branca, A., & Giraud, T. (2015). Adaptive horizontal gene transfers between multiple cheese-associated Fungi. Current Biology, 25, 2562–2569. 10.1016/j.cub.2015.08.025

Sato, Y., Bex, R., van den Berg, G. C. M., Santhanam, P., Höfte, M., Seidl, M. F., & Thomma, B. P. H. J. (2025). Starship giant transposons dominate plastic genomic regions in a fungal plant pathogen and drive virulence evolution. Nature Communications, 16(1), 6806. 10.1038/s41467-025-61986-6

Skotte, L., Korneliussen, T. S., & Albrechtsen, A. (2013). Estimating Individual Admixture Proportions from Next Generation Sequencing Data. Genetics, 195(3), 693–702. 10.1534/genetics.113.154138

Song, Y., Liu, X., Zhao, K., Ma, R., Wu, W., Zhang, Y., Duan, L., Li, X., Xu, H., Cheng, M., Qin, B., & Qi, Z. (2024). A new endophytic Penicillium oxalicum with aphicidal activity and its infection mechanism. Pest Management Science, 80(11), 5706–5717. 10.1002/ps.8288

Urquhart, Vogan, A. A., Gardiner, D. M., & Idnurm, A. (2023). Starships are active eukaryotic transposable elements mobilized by a new family of tyrosine recombinases. Proceedings of the National Academy of Sciences, 120(15), e2214521120. 10.1073/pnas.2214521120

Urquhart, Vogan, A. A., & Gluck-Thaler, E. (2024). Starships: A new frontier for fungal biology. *Trends in Genetics*, S0168952524001835. 10.1016/j.tig.2024.08.006

Van Dongen, S. (2008). Graph Clustering Via a Discrete Uncoupling Process. SIAM Journal on Matrix Analysis and Applications, 30(1), 121–141. 10.1137/040608635

van Kempen, M., Kim, S. S., Tumescheit, C., Mirdita, M., Lee, J., Gilchrist, C. L. M., Söding, J., & Steinegger, M. (2024). Fast and accurate protein structure search with Foldseek. Nature Biotechnology, 42(2), 243–246. 10.1038/s41587-023-01773-0

Xie, X.-H., Fu, X., Yan, X.-Y., Peng, W.-F., & Kang, L.-X. (2021). A Broad-Specificity Chitinase from Penicillium oxalicum k10 Exhibits Antifungal Activity and Biodegradation Properties of Chitin. Marine Drugs, 19(7),356. 10.3390/md19070356

Zhu, P., Weber, B., Rosenkranz, M., Ghirardo, A., & Schnitzler, J.-P. (2026). Volatile cues from pathogenic, mutualistic and saprotrophic fungi cause specific, fungus-dependent responses in Poplar. *Tree Physiology*, tpag053. 10.1093/treephys/tpag053

